# Optogenetic BMP4 models symmetry breaking and primitive streak formation in hPSCs

**DOI:** 10.64898/2026.10.01.755984

**Authors:** Alena Y. Streletskaia, Caleb J. Bashor, Aryeh Warmflash

**Affiliations:** Department of Biosciences, Rice University, Houston, TX 77030, USA; Department of Bioengineering, Rice University, Houston, TX 77030, USA; Ken Kennedy Institute, Rice University, Houston, TX 77005, USA; Rice Synthetic Biology Institute, Houston, TX 77005, USA; Division of Developmental Biology, Cincinnati Children’s Hospital Medical Center, Cincinnati, OH 45229, USA; Center for Stem Cell and Organoid Medicine, Cincinnati Children’s Hospital Medical Center, Cincinnati, OH 45229, USA; Department of Pediatrics, University of Cincinnati College of Medicine, Cincinnati, OH 45267, USA

## Abstract

Gastrulation begins when the epiblast epithelium breaks symmetry to form the primitive streak, but current in vitro models neither fully capture this process nor separate its mechanical and chemical inputs. Here, we introduce LightBMP, an optogenetic platform that enables spatiotemporally programmable BMP4 expression in human pluripotent stem cells. Dose and duration control reveal a sustained low-BMP regime, inaccessible to recombinant ligand, that specifies mesoderm. Spatially programmed BMP4 sources in uniform cultures mimic the radial organization of 2D gastruloids, showing that mechanical context and edge sensing are not required for patterning. In micropatterned colonies, light-defined sources break symmetry to generate a stripe-shaped, primitive-streak-like domain where cells acquire multiple mesodermal fates and undergo directed migration. Together, these results show that spatial control of a single morphogen source is sufficient to drive symmetry breaking and primitive-streak-like patterning, independent of edge constraints.

## INTRODUCTION

Gastrulation is a crucial stage of mammalian development during which pluripotent epiblast cells exit pluripotency and reorganize into the three primary germ layers: ectoderm, mesoderm, and endoderm.^1,2^ This transformation is orchestrated by morphogen signals whose spatial distribution, temporal dynamics, and dosage instruct region-specific fate decisions and coordinate large-scale tissue rearrangements.^3,4^ Among these signals, Bone Morphogenetic Protein 4 (BMP) plays a pivotal regulatory role. At the onset of gastrulation, BMP is expressed in extraembryonic tissues and induces primitive streak formation at the posterior of the epiblast.^5,6^ As the primitive streak elongates distally, BMP signaling extends to the proximal-posterior epiblast, and mesodermal cells that ingress early or via the posterior regions of the primitive streak adopt more lateral mesodermal identities.^7–10^ Genetic disruption of BMP signaling components results in failure of primitive streak formation or abnormal mesodermal patterning.^11–15^ Despite its essential role, how spatial and temporal patterns of BMP activity direct cell fate decisions and morphogenesis remains poorly understood, particularly in human development.

Direct manipulation of morphogen dynamics in mammalian embryos remains technically challenging and is not feasible in humans. Human pluripotent stem cells (hPSCs) provide an experimentally tractable platform for modeling human development;^1,16^ however, current systems do not recapitulate symmetry breaking in the context of an epithelial monolayer. In standard cultures, uniformly applied recombinant BMP produces relatively homogeneous differentiation.^17–19^ In geometrically confined 2D gastruloids, BMP induces self-organized concentric fate domains,^20–23^ while the radial symmetry of the colony is preserved. Asymmetric fate organization has been observed in 3D aggregates and embryo-like models,^24–27^ but these systems are difficult to manipulate quantitatively and do not reproduce the monolayer epithelial geometry of the pre-gastrulation epiblast. Recent optogenetic approaches have enabled spatially resolved induction of BMP expression;^28^ however, these approaches have not been used to generate asymmetric BMP sources capable of initiating streak formation. Together, these limitations have left open fundamental questions about how BMP dynamics initiate symmetry breaking and organize the body plan in human development.

To overcome this barrier, we introduce LightBMP, an optogenetic platform that enables tunable, reversible, and spatially and temporally programmable BMP expression. Using this system in hPSCs, we uncovered a previously inaccessible signaling regime: in contrast to the transient signaling elicited by comparable doses of recombinant BMP, LightBMP sustains low-level signaling that robustly induces mesodermal differentiation. Leveraging spatial control, we show that spatially patterned LightBMP activation generates graded BMP signaling that organizes mesodermal cells in standard culture into territories of lateral, paraxial, and axial subtypes reminiscent of the mesodermal organization of the embryo. Light-induced BMP expression generates fate patterns that diverge from the colony geometry, demonstrating that symmetry breaking is determined by BMP availability rather than by physical edge-sensing cues. Finally, by locally inducing BMP, we recapitulated core features of the human primitive streak *in vitro*, including localized epithelial-to-mesenchymal transition (EMT) and directed cell migration.

## RESULTS

### LightBMP: an optogenetic platform for controllable BMP secretion

To achieve spatiotemporal control of BMP4 (BMP) secretion in human pluripotent stem cells (hPSCs), we developed an optogenetic circuit comprising the blue-light–inducible transcription factor GAVPO^29^ and the human BMP4 precursor (proBMP) (**Fig. 1A**). GAVPO consists of a truncated Gal4 DNA-binding domain fused to the light-sensitive Vivid protein and p65 activation domain, with a C-terminal FLAG-tag for detection (**Fig. S1A**). Blue-light illumination induces GAVPO dimerization, which promotes transcription of a cassette containing a near-infrared fluorescent reporter (iRFP670)^30^ and proBMP, separated by a “self-cleaving” P2A^31^ peptide. proBMP is processed through the canonical secretory pathway and released as a mature BMP.^32,33^ BMP signaling activity in recipient cells is detected through the nuclear accumulation of phosphorylated SMAD1/5/8 (**Fig. 1A**).^34^

**Figure 1.**
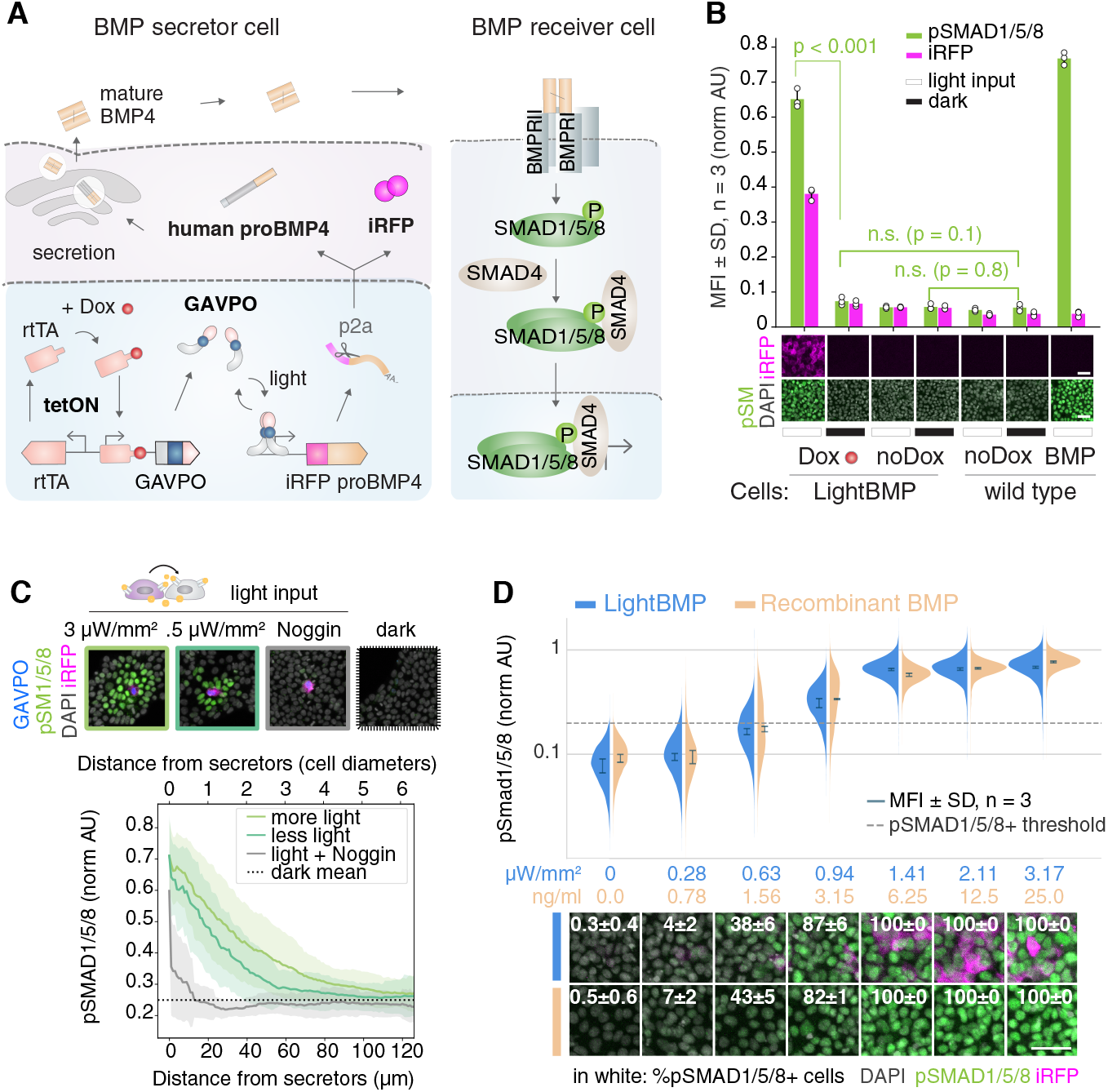
LightBMP: controllable morphogen secretion in human pluripotent stem cells (hPSCs). **(A)** Schematic of the LightBMP optogenetic circuit. Upon doxycycline (Dox) and blue-light exposure, the GAVPO transcription factor is expressed and dimerizes to activate a transgene encoding the near-infrared reporter iRFP670 (iRFP) and human BMP4 precursor (proBMP4), separated by a P2A self-cleaving peptide. iRFP marks BMP4-secreting cells; proBMP4 is processed through the canonical secretory pathway and secreted as mature BMP4 (BMP). BMP signaling activity in recipient cells is detected by immunostaining for phosphorylated SMAD1/5/8 (pSMAD1/5/8). **(B)** LightBMP hPSC line shows robust pSMAD1/5/8 activation upon Dox and blue light stimulation for 24 hours, while the background signaling is comparable to that of uninduced wild-type control cells. Scale bar, 50 μm. n = 3 wells per condition with 4 images per well, p < 0.001 by Student’s t-test. AU, arbitrary units; MFI, mean fluorescence intensity; SD, standard deviation. **(C)** Paracrine BMP signaling from LightBMP secretor cells. LightBMP cells (∼1% of total; identified by GAVPO-FLAG immunostaining and iRFP expression) were mixed with wild-type hPSCs and stimulated with blue light. pSMAD1/5/8 gradients extending up to 5–6 cell diameters beyond iRFP^+^ cells confirm paracrine BMP secretion. The gradient range is reduced at lower light intensity and abolished by NOGGIN co-treatment. Scale bar, 50 μm. n = 9 images per condition (∼20 LightBMP cells per image). **(D)** Dose-dependent BMP signaling output from LightBMP cells is comparable to recombinant BMP. Varying light doses (10 s pulses every 100 s for 24 h) produce graded pSMAD1/5/8 responses comparable to those elicited by titrated recombinant BMP. Scale bar, 50 μm. n = 3 wells per condition with 4 images per well.

To validate and optimize circuit performance, GAVPO and proBMP constructs were integrated into ESI-017 hPSCs using ePiggyBac transposition (**Fig. S1A**).^35^ Bulk populations were enriched for transgene-containing cells by antibiotic selection and analyzed by immunofluorescence (**Fig. S1B–D**). Compared with the other promoters tested, GAVPO expression driven by the strong doxycycline-inducible TRE promoter yielded robust light-dependent induction of iRFP and accumulation of pSMAD1/5/8, suggesting that sufficient GAVPO abundance is required for circuit activation (**Fig. S1B**). Importantly, GAVPO protein levels remained stable throughout 10 days of continuous doxycycline exposure, suggesting resistance to transgene silencing under prolonged induction (**Fig. S1C**). Because leaky BMP expression could induce unwanted differentiation of hPSCs, tighter control of basal activity was required. To address this, we compared three promoter variants driving iRFP-P2A-proBMP by quantifying iRFP fluorescence as a readout of promoter activity (**Fig. S1D**). The variant with a weaker minimal promoter and fewer GAVPO-binding sites reduced background iRFP expression in the dark (**Fig. S1D**).

To ensure uniform light responsiveness, we single-cell–sorted the bulk population, screened 21 clonal lines under light and dark conditions, and selected a clone exhibiting minimal basal activity (**Fig. S1E**). The LightBMP cell line showed a robust increase in iRFP expression and pSMAD1/5/8 signaling after 24 hours of light stimulation, with no significant difference from wild-type cells in the dark or in the absence of doxycycline (**Fig. 1B**). However, we observed a small but measurable degree of spontaneous differentiation during routine passaging and evaluated whether culture with BMP inhibitors would preserve pluripotency (**Fig. S1F**). Prolonged treatment with the BMPR1 kinase inhibitor LDN193189^36^ maintained cells in pluripotency but compromised subsequent differentiation into ISL1⁺ amnion after withdrawal (**Fig. S1G**). In contrast, long-term culture with recombinant NOGGIN preserved both pluripotency, as judged by cell morphology and the expression of OCT4 (POU5F1), SOX2, and NANOG, as well as the ability of cells to differentiate in response to recombinant BMP at levels comparable to those of the wild-type control (**Fig. S1F-I**).

To confirm that LightBMP cells can signal to their neighbors in a paracrine manner, we separated “source” and “sink” populations by mixing a small fraction (∼1%) of LightBMP cells with wild-type cells (**Fig. 1C**). Following blue light stimulation, LightBMP cells (identified by FLAG-tagged GAVPO immunostaining and iRFP expression) generated pSMAD1/5/8 signaling gradients extending up to 5–6 cell diameters into the surrounding wild-type population, indicating secretion and sensing by neighboring cells (**Fig. 1C**). Gradient range was reduced at a lower light intensity and was abolished by NOGGIN treatment (**Fig. 1C**).

To estimate how much BMP is secreted by LightBMP hPSCs, we compared signaling responses elicited by graded light intensities to those induced by defined concentrations of recombinant BMP, assessing both conditions at 24 hours (**Fig. 1D**). Light stimulation produced a dose-dependent increase in pSMAD1/5/8 activity that closely recapitulated the response curve obtained with recombinant BMP (**Fig. 1D**). Together, these results establish LightBMP hPSCs as a tunable optogenetic platform for dose-controlled BMP secretion.

### LightBMP provides quantitative, tunable control over BMP signaling

To systematically test LightBMP responses across illumination regimes, we built a custom high-throughput LED device, optoBox (**Fig. 2A**). Inspired by previously reported hardware,^37–40^ optoBox allows independent light programming in each well, enabling 96 parallel stimulation conditions (**Fig. 2A, Fig. S2A**). A distinguishing feature of optoBox is the placement of a visible-light bandpass filter between each LED and the well bottom, which reduces sample heating while preserving a linear relationship between programmed and delivered light (**Fig. S2A-C**). This setup allows reliable, high-throughput optogenetic experiments in a tissue culture incubator, with minimal cell death at light doses below 3 × 10⁵ µJ/mm² (**Fig. 2A**; **Fig. S2D**).

**Figure 2.**
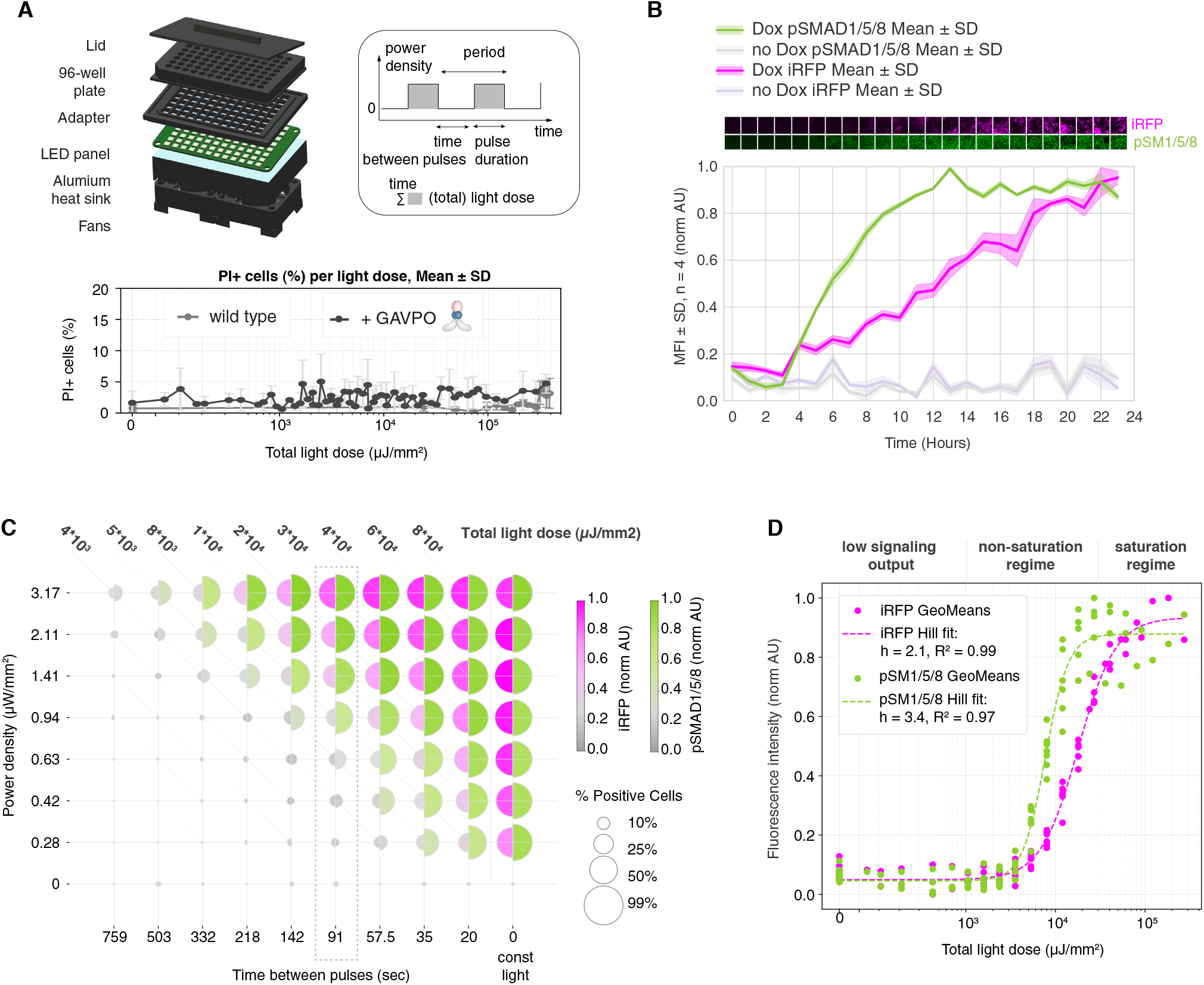
LightBMP provides quantitative, tunable control over BMP signaling. **(A)** The optoBox high-throughput illumination device enables independent light programming across a 96-well plate. Top left: schematic of device assembly. Top right: diagram of light-dose parameters: power density (intensity), pulse duration, and inter-pulse interval together determine total light dose. Bottom: propidium iodide (PI)-positive cell percentage as a function of total light dose, showing that light doses below 3 × 10^5^ µJ/mm^2^ do not compromise viability in LightBMP (black) or wild-type (gray) hPSCs. n = 3 images per condition. **(B)** Kinetics of BMP signaling under continuous illumination (0.8 µW/mm^2^). Time course of iRFP and pSMAD1/5/8 mean fluorescence intensities in doxycycline-treated (Dox) versus doxycycline-untreated (no Dox) LightBMP cells. pSMAD1/5/8 is detectable by 4 h and reaches a plateau by ∼12 h; iRFP continues to increase over 24 h. Data are mean ± SD; n = 4 images per condition. **(C)** Systematic mapping of pSMAD1/5/8 responses to constant and pulsed light regimes. Each heatmap point represents a distinct light condition, with a fixed pulse duration of 10 s and varying power density (intensity) and pulse frequency. Higher intensity and shorter intervals between pulses increase BMP output. Diagonal bands correspond to equivalent total light doses. Geometric means of ∼4 × 10³ cells per condition are shown for fluorescence intensity immunostaining. **(D)** The relationship between total light dose and steady-state pSMAD1/5/8 response follows a Hill function. n = 4 images per condition.

To define LightBMP activation kinetics and determine when signaling reaches a steady state, we quantified pSMAD1/5/8 under continuous (0.8 µW/mm², **Fig. 2B**) or pulsed (10-s pulses every 100 s) illumination (**Fig. S2E**). Under continuous illumination, both iRFP and pSMAD1/5/8 expression were detectable by 4 hours, with pSMAD1/5/8 expression plateauing by ∼12 hours and iRFP continuing to rise over the 24-hour window (**Fig. 2B**). Pulsed illumination induced pSMAD1/5/8 levels that plateaued between 16 and 24 hours, with the level depending on the illumination intensity (**Fig. S2E**). We therefore selected 24 hours as a consistent time point to assess a steady-state LightBMP output.

We measured pSMAD1/5/8 signaling under continuous and pulsed illumination, systematically varying light intensity and inter-pulse interval, while maintaining a fixed pulse duration of 10 s (**Fig. 2C**). Increasing either intensity or pulse frequency resulted in higher BMP signaling output (**Fig. 2C**). Notably, activation of only ∼30% of iRFP⁺ cells was sufficient to achieve a maximal signaling response, consistent with paracrine BMP action (**Fig. S2F**). Across conditions, light intensity and pulse frequency were functionally interchangeable, and the integrated light input, defined as total light dose, strongly predicted steady-state BMP signaling levels (**Fig. 2D**; R² = 0.97; **Fig. 2C** diagonals).

The relationship between total light dose and pSMAD1/5/8 signaling followed a Hill function (**Fig. 2D**), with the non-saturating region corresponding to signaling levels equivalent to low-dose recombinant BMP at the same tested conditions (**Fig. 1D**, **Fig. 2C** dashed column). However, the kinetics by which these steady-state levels are achieved differ between the two approaches: low-dose recombinant BMP elicits a strong initial response that decays as ligand is lost from the culture medium,^17–19,41^ whereas LightBMP induces a gradually increasing pSMAD1/5/8 response that plateaus over time (**Fig. 2B; Fig. S2E**). This behavior enables non-decaying low BMP signaling, a regime that is technically challenging to achieve using recombinant ligand alone.^19,41^ Similar low levels of steady-state signaling can be generated by using saturating doses of BMP and titrating the BMP receptor inhibitor LDN193189 to derive amnion-like cells; however, this strategy is not generally applicable to other differentiation contexts, as high concentrations of LDN193189 can affect downstream pathways due to limited receptor selectivity.^41,42^ In contrast, LightBMP sustains low-level signaling through continuous ligand production, as pSMAD1/5/8 levels decline rapidly when light is turned off, particularly after low-intensity exposure (**Fig. S2E**). Together, these results establish LightBMP as a reversible, light-tunable system for dynamic and dose-controlled manipulation of BMP signaling.

### Light-induced low BMP establishes a distinct route to mesoderm

Previous work from our group demonstrated that BMP signaling duration determines cell fate: short duration maintains pluripotency marked by SOX2, intermediate duration induces mesoderm-like differentiation marked by Brachyury (BRA, or TBXT), and prolonged signaling activates amnion genes such as ISL1 (**Fig. 3A**).^19^ Using these established outcomes as a reference, we tested how LightBMP dynamics direct cell fate decisions. LightBMP hPSCs were stimulated for durations ranging from 6 to 48 h at light intensities between 0.28 and 3.17 µW/mm² (**Fig. 3B**). Illumination was delivered as 10-s pulses every 100 s to produce a diverse range of steady-state signaling outputs (**Fig. 2C** dashed column). Cells were maintained in ROCK inhibitor, following our previously reported experiments with recombinant BMP, which improves response uniformity, and fixed for immunostaining at 48 h, a time point sufficient to capture differentiation outcomes.^19^

**Figure 3.**
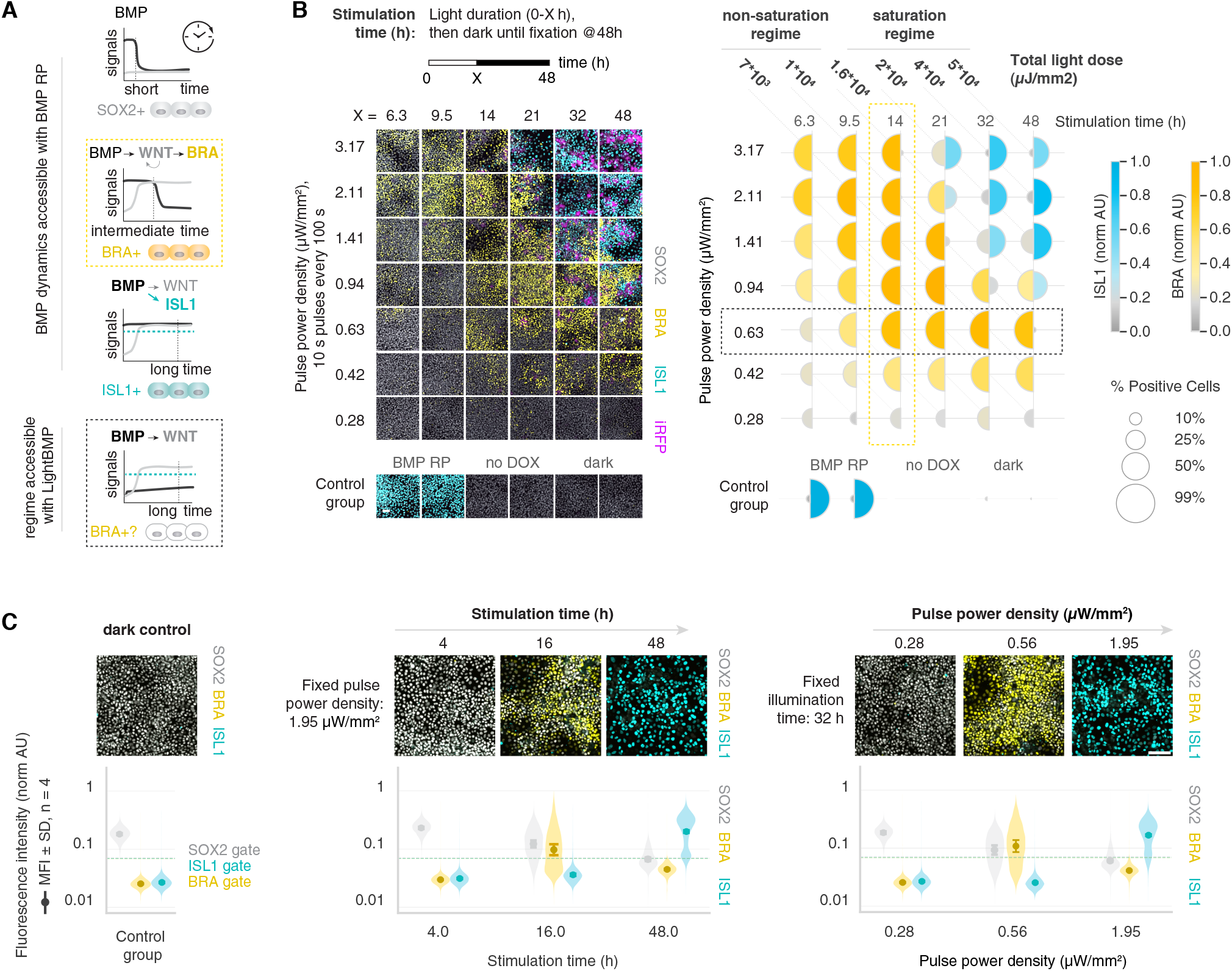
Light-induced low BMP establishes a distinct route to mesoderm. **(A)** Schematic of BMP signaling dynamics and cell fate outcomes based on previous findings with recombinant BMP: short exposure maintains pluripotency (SOX2^+^); intermediate duration induces mesoderm (BRA^+^; yellow dashed rectangle); prolonged exposure promotes amnion fate (ISL1^+^).¹⁹ Low, sustained BMP sufficient to activate WNT but below the ISL1 threshold was hypothesized to promote BRA⁺ mesoderm (black dashed rectangle). **(B)** LightBMP stimulation recapitulates duration-dependent fate responses at high light intensity, with BRA expression peaking at intermediate durations (yellow dashed rectangle) and prolonged exposure favoring ISL1. Lower light intensity robustly induces BRA without ISL1 even after 48 h, defining a sustained low-level BMP regime for mesoderm induction (black dashed rectangle). Scale bar, 50 μm. Geometric means of ∼6 × 10³ cells per condition shown for fluorescence intensity immunostaining. **(C)** Demonstration of dual control mechanisms for gastrulation-like fate specification via illumination duration (time) and light intensity (dose). Scale bar, 100 μm. n = 16 images per condition.

At high light intensities (1.41, 2.11, and 3.17 µW/mm²), LightBMP recapitulated duration-dependent BMP responses: shorter stimulation (<21 h) promoted BRA, whereas prolonged stimulation (>21 h) induced ISL1 expression (**Fig. 3B**; **Fig. S3A**). ISL1 upregulation occurred earlier at higher intensities and at higher cumulative light doses (2–5 × 10⁴ µJ/mm²), consistent with previous reports indicating that amnion-like differentiation is regulated by the time-integral of BMP signaling input.^41^ Low light doses (7 × 10³ and 1 × 10⁴ µJ/mm²) favored mesodermal differentiation (**Fig. 3B**). Unlike the high-intensity regime, where BRA expression peaked at shorter light durations, low intensities (0.42 and 0.63 µW/mm²) maintained BRA expression even after 48 h of stimulation (**Fig. 3B**; **Fig. S3A**). A similar accumulation of BRA⁺ cells was observed in the absence of ROCK inhibitor, although with substantially fewer mesodermal cells (**Fig. S3B**).

High BRA expression after prolonged stimulation contrasts with recombinant 48-h BMP treatment, which does not efficiently induce mesoderm at any dose (**Fig. S1I**).^19^ At low intensities, LightBMP produces low, sustained BMP signaling that may be sufficient to induce mesoderm-like differentiation but insufficient to activate ISL1 (**Fig. 3A**, bottom). While BMP signaling can directly induce ISL1, BRA induction occurs through the downstream morphogen WNT3 (hereafter WNT) (**Fig. 3A**).^19,22,41,43^ To assess WNT pathway activity under these conditions, we immunostained for LEF1, a direct WNT target and nuclear effector of WNT.^44,45^ LEF1 was upregulated in both BRA-high and ISL1-high conditions, indicating that BMP signaling activates the WNT pathway in both differentiation regimes and suggesting that WNT activity alone might be insufficient to distinguish between mesoderm-like and amnion-like cell fates (**Fig. S3C**). We therefore examined whether differences in BMP signaling might account for these outcomes. pSMAD1/5/8 activity could not be directly compared across different stimulation durations because BMP signaling decays rapidly upon light removal (**Fig. S2E**). However, under identical stimulation durations, higher pSMAD1/5/8 activity was associated with increased ISL1 expression (**Fig. S3C**). These observations suggest that mesodermal differentiation occurs at BMP signaling levels sufficient to sustain WNT activity but below the threshold required for ISL1 induction, thus paralleling the duration-dependent BMP model proposed by Camacho-Aguilar et al.^19^

Consistent with the observed interplay between illumination duration and intensity, pluripotency markers (SOX2 and NANOG) were preserved under brief or low-intensity illumination, intermediate stimulation promoted mesodermal marker expression (BRA and MIXL1), and prolonged or high-intensity illumination drove amnion-like fates (ISL1 and GATA3) (**Fig. 3C**; **Fig. S3D**). These results show that differentiation outcomes can be precisely tuned by modulating either illumination duration or intensity, with mesodermal differentiation preferentially emerging at lower light doses than amnion-like cells.

### Optogenetically-defined BMP gradients drive spatially ordered mesodermal patterning

To test the ability of the LightBMP system to generate spatial gradients of BMP ligands, and to probe the signaling and differentiation response to them, we illuminated one half of a uniform hPSC culture to induce BMP production while the adjacent half remained protected from light. iRFP-P2A-proBMP expression was restricted to the illuminated secretor region, forming a sharp iRFP⁺ boundary with the neighboring receiver cells (**Fig. 4**, **Fig. S4A**). Cultures were fixed following days 2, 3, and 4 of illumination (**Fig. 4A**, **Fig. S4A**).

**Figure 4.**
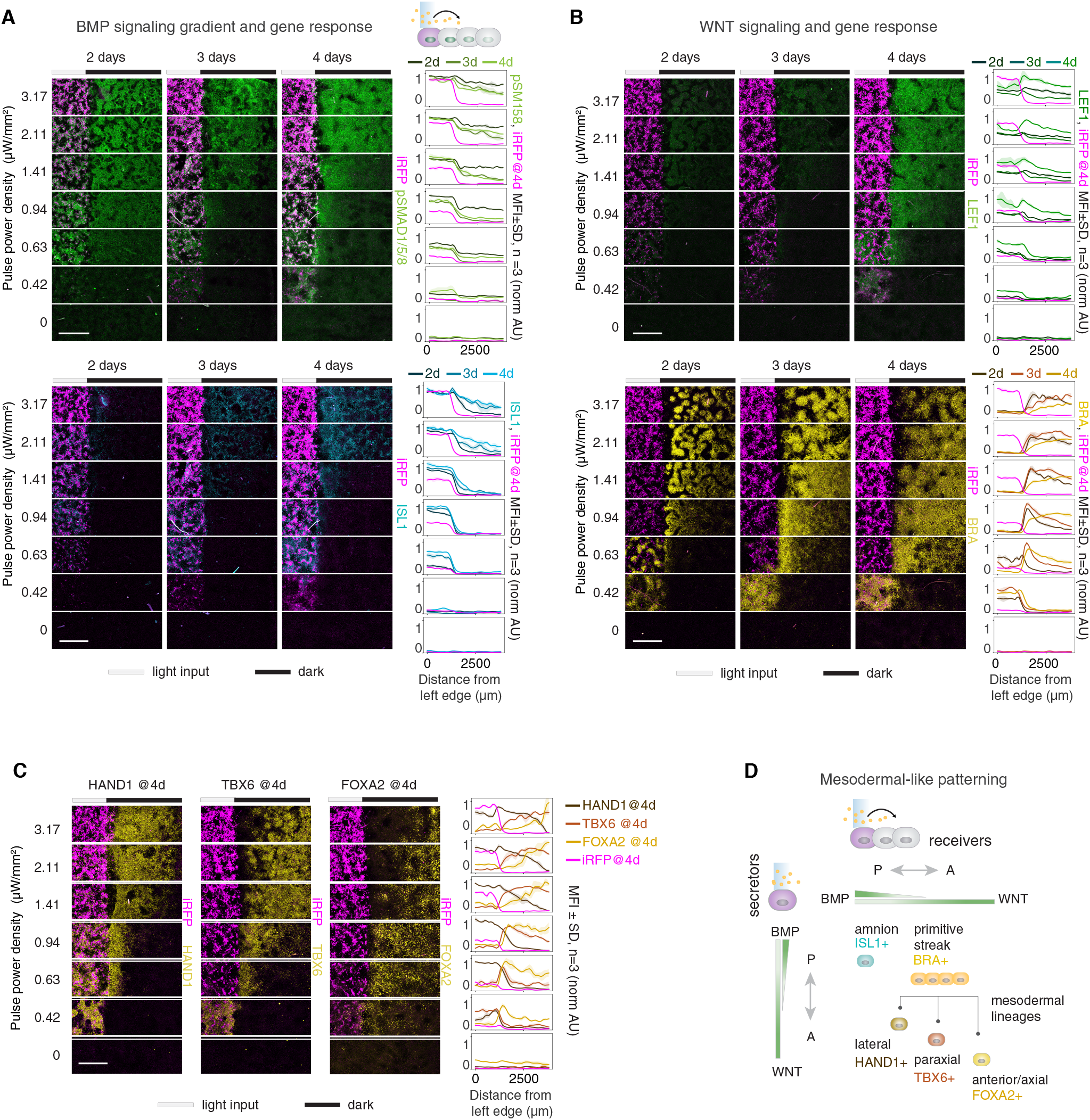
Optogenetically-defined BMP gradients drive spatially ordered mesodermal pat-terning. **(A)** Spatial BMP gradients and ISL1 induction in half-illuminated cultures. One half of each well (iRFP^+^ secretor region) was illuminated for 2, 3, or 4 days; the adjacent half remained dark (receiver region). Immunostaining for pSMAD1/5/8 and ISL1 is shown at each time point and stimulation intensity. Line graphs show mean fluorescence intensity profiles ± SD along the gradient axis. Scale bar, 1000 μm. n = 3 image grids (1 × 3 fields) per condition. **(B)** Downstream responses to BMP gradients. Immunostaining for LEF1 (WNT signaling target) and BRA at days 2, 3, and 4. Scale bar, 1000 μm. n = 3 image grids (1 × 3 fields) per condition. **(C)** Spatial organization of mesodermal transcription factors along the BMP gradient at day 4. HAND1 (lateral plate mesoderm), TBX6 (paraxial mesoderm), and FOXA2 (axial mesoderm) are expressed at progressively greater distances from the BMP source. Scale bar, 1000 μm. n = 3 image grids (1 × 3 fields) per condition. **(D)** Schematic summarizing BMP gradient-driven patterning. Optogenetically defined BMP gradi-ents generate a continuum of mesodermal identities corresponding to lateral plate- (HAND1⁺), paraxial- (TBX6⁺), and axial- (FOXA2⁺) mesoderm-like fates.

Illumination intensity set both the amplitude and the spatial range of the resulting signaling gradient. Higher light intensities produced pSMAD1/5/8 gradients that extended over longer ranges (**Fig. 4A**). For a given intensity, pSMAD1/5/8 levels peaked on day 2 and then declined to a steady level on days 3 and 4 (**Fig. 4A**). The drop following day 2 likely reflects the removal of ROCK inhibitor following the first 2 days of the protocol, as this molecule improves the accessibility of BMP receptor complexes.^21,46,47^ Consistent with this, cultures maintained in ROCK inhibitor showed elevated pSMAD1/5/8 extending further into non-illuminated regions than cultures where ROCK inhibitor was withdrawn before illumination (**Fig. S4B**).

As above, cells with the highest BMP signaling, primarily those in the illuminated region, induced ISL1 expression by day 2, whereas cells experiencing lower levels of signaling did not (**Fig. 4A**; **Fig. S3B-C**). Cells receiving insufficient BMP doses to convert to amnion fates may adopt mesodermal fates due to induced WNT signaling, so we stained for the direct WNT target LEF1 as well as for BRA to mark primitive streak cells transitioning to mesoderm.^44,45^ In contrast to the pSMAD1/5/8 gradients, which had stabilized by day 3, LEF1 gradients increased in amplitude and extended over longer ranges by day 4 (**Fig. 4B**). Over the same interval, receiver cells that had not reached the ISL1 “threshold” induced BRA, and by day 4 BRA⁺ cells extended hundreds of micrometers from the light-dark boundary (**Fig. 4B**). The correspondence between the expanding LEF1 and BRA⁺ domains is consistent with WNT acting downstream of the BMP signal to specify primitive streak identity in cells below the amnion threshold.

Within this BRA⁺ field, cells progressively acquired distinct mesodermal identities ordered by their distance from the BMP source. By day 4, cells closest to the source preferentially expressed the lateral plate mesoderm marker HAND1, followed by a paraxial-like TBX6-expressing region at intermediate distances and an axial-like FOXA2-expressing domain furthest from the source (**Fig. 4C**).^48,49^ Separation between HAND1⁺ and TBX6⁺ regions was already evident at day 3 (**Fig. S4C**) and progressed further by day 4 as the BRA⁺ domain continued to expand (**Fig. 4B**). The domains were arranged sequentially along the gradient with gradual boundaries, reflecting a transcriptional continuum rather than discrete fates.

Additional lineage markers assayed at day 4 supported these assignments (**Fig. S4D**). FOXF1 largely overlapped with HAND1⁺ cells and extended partially into the TBX6⁺ region, consistent with lateral mesoderm identity.^50^ Sparse NKX2.5⁺ cells were detected mainly within light-exposed regions overlapping the ISL1⁺ and HAND1⁺ domains, suggesting the presence of heart-forming progenitors.^48,49^ FOXC2 appeared at low, patchy levels within the TBX6⁺ domain, potentially reflecting maturation toward paraxial mesoderm or initiation of anterior somite-like programs.^48,51^ The majority of FOXA2⁺ cells did not express SOX17, consistent with their identity as axial mesoderm rather than endoderm.^48,49^

The resulting arrangement of lateral plate–, paraxial–, and axial mesoderm–like domains parallels observations in mouse embryos, where mesodermal subtypes are specified along the medial-lateral axis in response to BMP signaling (**Fig. 4D**).^9,11,14,52,53^ Optogenetically defined BMP gradients are thus sufficient to break the symmetry of a uniform hPSC culture and to drive progressive, spatially ordered patterning of mesodermal fates in an embryo-like manner.

### Spatiotemporal BMP availability rather than geometric constraints determines fate patterning in micropatterned hPSCs

Micropatterned hESCs treated with recombinant BMP form radially symmetric patterns of gastrulation-associated cell fates, with amnion-like cells at the edge, an inner ring of mesoderm-like cells, and a pluripotent or ectoderm-like center.^20,54^ During this process, BMP signaling becomes spatially restricted to the colony edge, driven by the apical localization of BMP receptors in peripheral cells and by BMP inhibitor secretion from the colony center (**Fig. 5A**, top).^21,55^ As a result, the cells effectively “sense” their distance from the edge, leading to the loss of central fates in smaller colonies.^20^ We therefore asked whether these edge-sensing mechanisms are intrinsically required for differentiation or merely serve as a method to introduce graded BMP activity from ligand introduced into the bulk medium.

**Figure 5.**
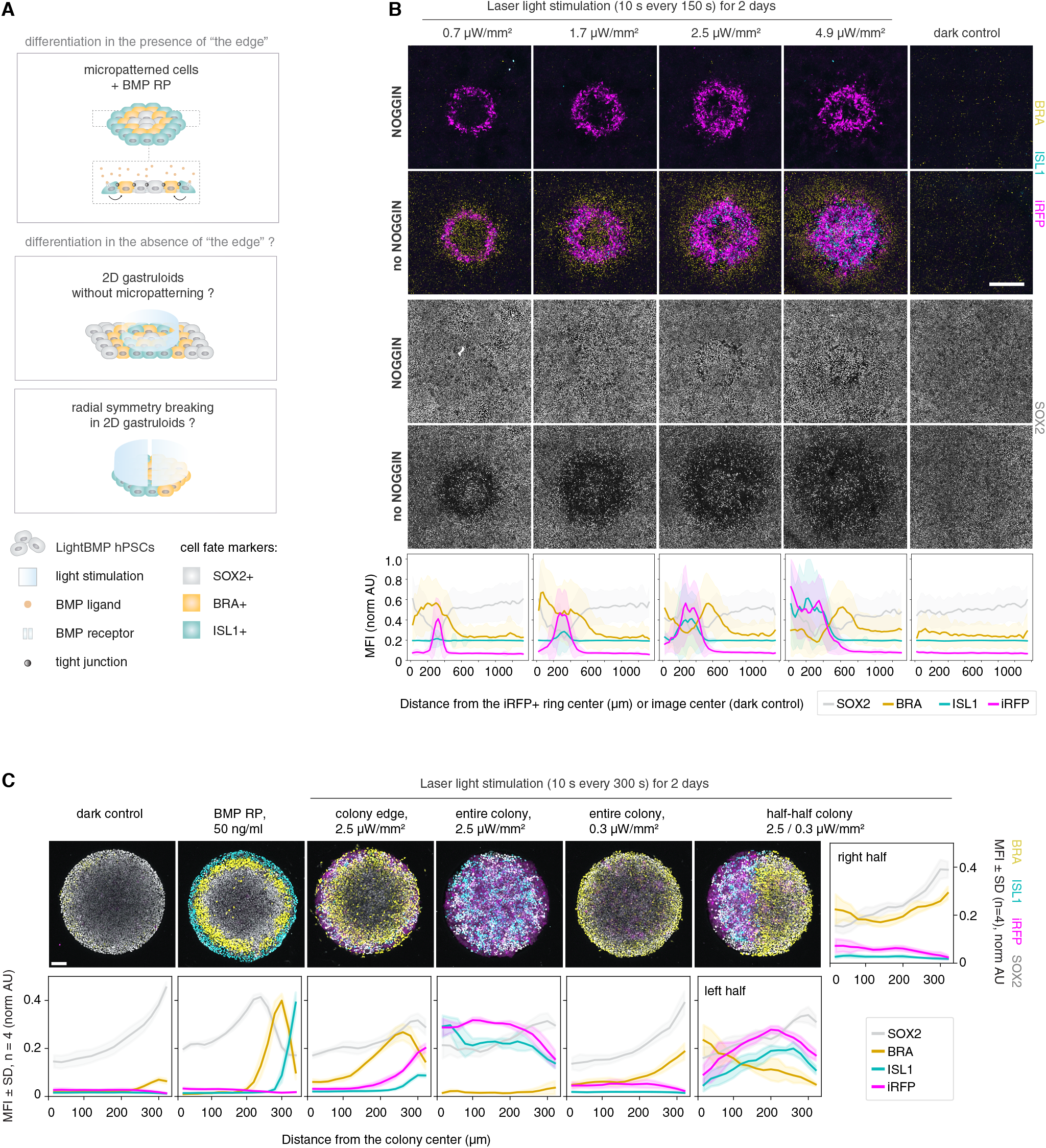
Spatiotemporal BMP availability rather than geometric constraints determines fate patterning in micropatterned hPSCs. **(A)** Schematic of BMP patterning in micropatterned colonies. Top: recombinant BMP induces edge-restricted signaling and radial fate organization due to receptor localization and inhibitor gradients. Middle (related to Fig 5B): optogenetically induced ring-shaped BMP secretion generates gastrulation-like radial patterns in standard culture without colony edges. Bottom (related to Fig 5C): LightBMP enables spatial control of differentiation independent of colony edges. **(B)** Ring-shaped laser stimulation for 2 days generates gastrulation-like fate patterns in uniform hPSC cultures lacking colony edges. Immunostaining at day 3 shows concentric ISL1^+^, BRA^+^, and SOX2^+^ domains around iRFP^+^ BMP-secretor rings, analogous to micropatterned 2D gastruloid organization. At the highest intensity (4.9 µW/mm^2^), an inverse ISL1–BRA–SOX2 order emerges from the ring periphery inward. Fluorescence profiles show mean intensity ± SD across cells vs. distance from the iRFP+ ring center (or image center for dark control). Scale bar, 500 μm. Note: only one ring could be generated per well under this stimulation regime; statistical quantification across multiple rings is presented in Figure S5. **(C)** Spatial control of differentiation in micropatterned colonies. Uniform illumination drives homogeneous fates across entire colonies, with high light inducing ISL1⁺ and lower light inducing BRA⁺ identities. Differential illumination across a single colony generates asymmetric patterning, with distinct ISL1⁺ and BRA⁺ domains on opposite sides. Residual edge effects are detectable but do not dominate overall patterning. Scale bar, 100 μm. n = 4 colonies per condition.

To test whether gastrulation-like differentiation can occur in 2D gastruloid-like patterns independently of colony edge constraints, ring-shaped BMP secretion was induced in uniformly seeded cultures lacking imposed differences in receptor distribution. Light stimulation was delivered using a laser-scanning microscope with a moving laser spot, applying a 10-s scan per ring repeated approximately every 150 s, comparable to the 10-s pulses delivered every 100 s in previous experiments (**Fig. 5B**). Alternatively, a lower-frequency regime (10-s per ring every ∼450 s) was used, reducing BMP output but allowing multiple rings to be generated per stimulation cycle and enabling statistical quantification across light conditions (**Fig. S5A–B**). At 2 days of stimulation, iRFP⁺ rings were observed with surrounding pSMAD1/5/8 signaling, suggesting successful patterning of BMP secretors at ∼100 μm resolution, as measured by ring width (outer minus inner diameter) (**Fig. S5A**).

By day 3, light-patterned BMP secretion had generated gastrulation-like fate responses around the illuminated rings. Cells within the iRFP⁺ rings adopted an amniotic-like identity marked by ISL1, while adjacent cells expressed BRA, followed by SOX2⁺ undifferentiated cells, with the inner SOX2⁺ region better preserved at lower light doses (**Fig. 5B**, **Fig. S5B**). Thus, the canonical ISL1-BRA-SOX2 ordering seen in micropatterned colonies was recapitulated in standard cultures lacking physical colony edges, suggesting that edge cues are not essential for differentiation (**Fig. 5A**, middle).

Notably, at the highest light dose, an “inverse” micropattern emerged, with SOX2-BRA-ISL1 arranged from the periphery of the induced ring toward the center (**Fig. 5B**: 4.9 μW/mm²), indicating that the center-edge axis can be reversed without substantially altering cell fate outcomes. These results suggest that BMP receptor localization and mechanical edge constraints are not crucial for fate patterning per se, but instead determine where BMP is sensed, shaping the final spatial organization of cell fates.

These findings further indicate that differentiation can be induced in any region of the colony where BMP is secreted (**Fig. 5A**, bottom). To directly test whether pattern-forming mechanisms are overridden by programmed BMP inputs, we illuminated entire micropatterned colonies. Depending on light intensity, the entire colony could be driven toward a uniform ISL1⁺ fate at high stimulation or a BRA⁺ fate at lower stimulation (**Fig. 5C**, entire colony stimulations). Edge effects were still detectable, with slightly higher SOX2 and BRA levels at the periphery, likely reflecting the sensing of TGF-β and FGF ligands from the mTeSR medium;^21,22,56^ however, overall patterning was dictated by the programmed BMP input.

We next tested whether radial symmetry could be actively disrupted by imposing distinct BMP inputs on opposite sides of a single colony. The left half was illuminated at 2.5 µW/mm², whereas the right half received 0.3 µW/mm². This differential stimulation generated spatially segregated domains within individual colonies, with one half adopting an ISL1⁺ identity and the other a BRA⁺ identity (**Fig. 5C**). Together, these results demonstrate that spatially programmed BMP secretion directs gastrulation-like fate patterning independently of edge-imposed constraints, and that the radial symmetry of 2D gastruloid responses can be actively controlled or disrupted.

### LightBMP induces symmetry breaking and primitive streak-like patterning in micropatterned hPSCs

Radial symmetry in BMP-treated micropatterned hPSCs fails to recapitulate the geometry of the human gastrula, where the primitive streak forms as a stripe rather than concentric rings.^50,57–60^ We therefore tested whether spatially localizing BMP signaling within micropatterned colonies could create a primitive-streak-like domain that better mimics human developmental patterning (**Fig. 6A**).

**Figure 6.**
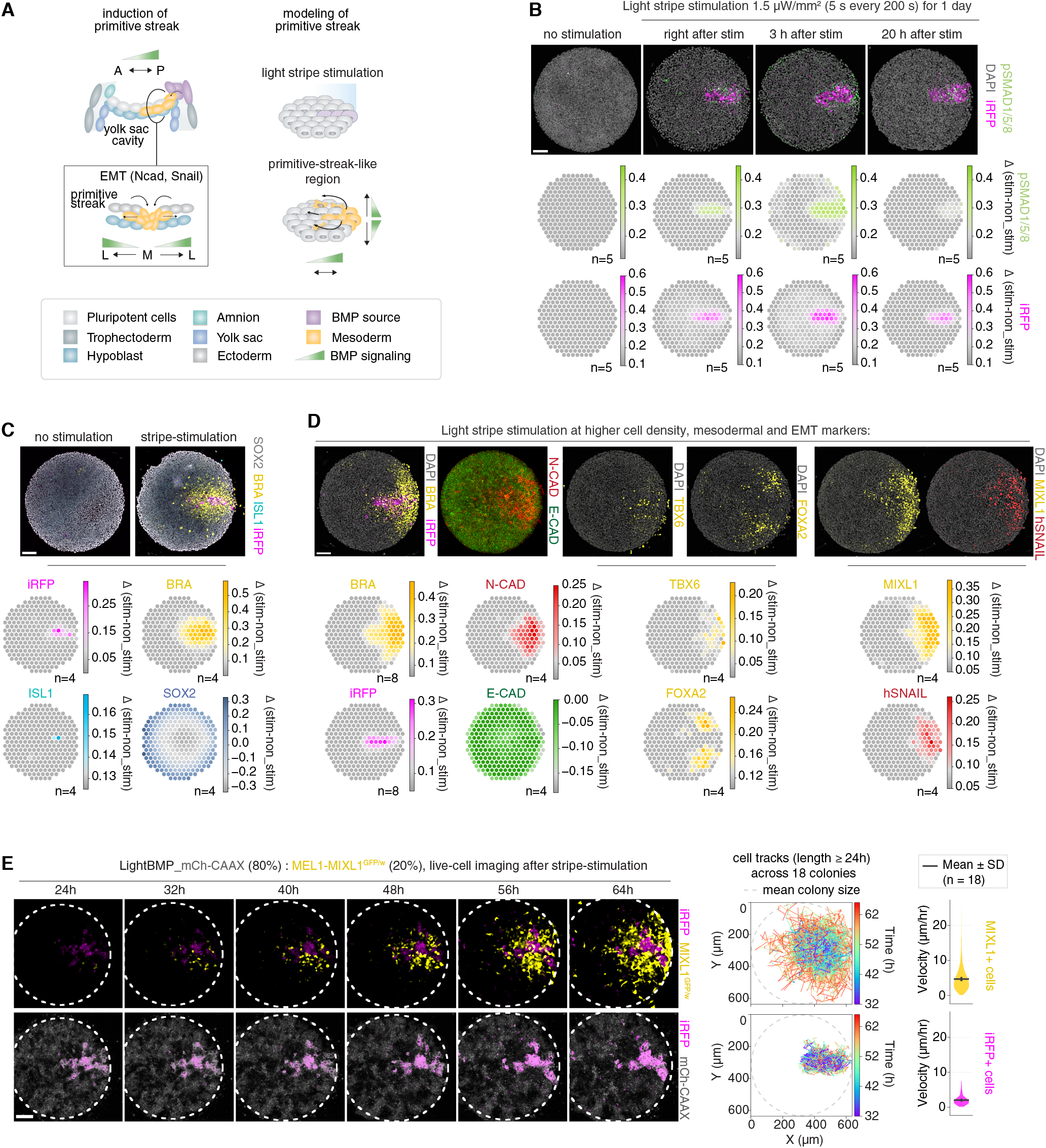
LightBMP induces symmetry breaking and primitive streak–like patterning in hPSCs. **(A)** Schematic of stripe-shaped laser stimulation to model primitive streak (PS) formation within micropatterned hPSC colonies. Low-dose illumination of a defined linear stripe at one side of the colony induces a localized BMP source predicted to break radial symmetry and generate a PS-like mesodermal domain. **(B)** Spatial confinement of BMP signaling. Stripe illumination induces iRFP⁺ BMP-producing cells restricted to the stimulated region. pSMAD1/5/8 signaling is confined to the stripe and immediate neighboring cells after stimulation and remains spatially localized following ROCK inhibitor withdrawal and during the light OFF phase. Scale bar, 100 μm. n = 5 colonies per condition. **(C)** Cell fate responses following stripe stimulation. At 48 h, low-dose BMP primarily induces BRA⁺ mesodermal fates, with rare ISL1⁺ cells restricted to regions of highest BMP output. BRA⁺ cells extend beyond the stimulated region, suggesting migration from the stripe-like domain. Scale bar, 100 μm. n = 4 colonies per condition. **(D)** Spatial organization of mesodermal and EMT markers. BRA⁺ cells expand from the stimulation region and overlap with MIXL1. TBX6 is enriched proximal to the BMP stripe, whereas FOXA2 appears in more distal regions, indicating anterior–posterior PS-like patterning. EMT-associated markers, including N-CAD and hSNAIL, are induced within the mesodermal domain. Scale bar, 100 μm. n = 4 or 8 colonies per condition. **(E)** Directed migration of primitive streak-like cells. LightBMP_mCherry-CAAX hPSCs (iRFP^+^/mCh-CAAX^+^; 80%) were co-cultured with MEL1-MIXL1^GFP/w^ reporter cells (20%) and subjected to stripe stimulation. MIXL1^+^ cells are induced near the iRFP^+^ stripe and migrate away from it. Left panels: representative time-lapse frames. Right: cell tracks (length ≥ 24 h) and velocity distributions for MIXL1^+^ and iRFP-high populations across 18 colonies. Scale bar, 100 μm. n = 18 colonies. See also Figure S6 and Video S1.

Mesoderm is efficiently induced under a low light-dose regime that produces BMP signaling sufficient for WNT propagation while remaining below the ISL1 activation threshold (**Figs. 3A–B, S3C**). Spatially restricted activation was achieved by delivering low-dose laser stimulation in a defined stripe pattern (**Fig. 6A**). Following stimulation, iRFP⁺ BMP-secretor cells were confined to the illuminated stripe, confirming spatially restricted activation (**Fig. 6B**). pSMAD1/5/8 immunostaining was found in the iRFP⁺ region and up to two cell diameters away, indicating low BMP output that activated signaling only in immediate neighbors (**Fig. 6B**, right after stimulation). ROCK inhibitor withdrawal after stimulation maintained the stripe-localized pSMAD1/5/8 distribution (**Fig. 6B**, 3 h post-stimulation). By 20 h, pSMAD1/5/8 levels decreased, consistent with LightBMP OFF responses at low doses (**Fig. S2E**), yet remained spatially confined to the illuminated iRFP-high region (**Fig. 6B**, 20 h post-stimulation). Together, these results demonstrate that patterned illumination breaks the symmetry of BMP signaling response and restricts pSMAD1/5/8 to a stripe-like domain within micropatterned colonies.

To verify predominantly mesodermal fate stimulation, we assessed cell fate responses at 48 hours. Rare ISL1⁺ cells were restricted to the iRFP-high region (strongest BMP secretion), while most cells in and around the illuminated domain expressed BRA (**Fig. 6C**). BRA⁺ cells were also detected far beyond the stimulated region, including outside colony boundaries, suggesting cell migration from the stripe-like region.

To examine primitive-streak-like features, we increased cell density to approximate high epiblast density^61^ and assessed additional mesodermal and EMT markers (**Fig. 6D**). BRA⁺ cells expanded toward colony edges near the stimulation region, overlapping with MIXL1, which was slightly displaced from the highest iRFP-marked BMP secretion. TBX6 was expressed proximal to the iRFP⁺ stripe, while FOXA2 appeared in more distal mesodermal regions, indicating spatially offset mesodermal populations with anterior–posterior primitive-streak-like organization.^48,49,62^

Epithelial-to-mesenchymal transition (EMT) markers N-CAD and hSNAIL were also expressed at the primitive-streak-like region, with N-CAD upregulated more broadly and hSNAIL more spatially restricted to the proximal region of the stripe (**Fig. 6D**).^63^ N-CAD induction is consistent with cadherin switching during EMT, although E-CAD remained detectable, likely reflecting the multilayered micropattern structure (2-3 cell layers), in which not all layers simultaneously downregulate E-CAD. hSNAIL, which represses E-CAD expression, was still observed (**Fig. 6D**).^64^

Primitive streak formation involves directed anterior–lateral migration of mesodermal cells.^65^ To test whether primitive-streak-like cells exhibited similar directional movement, LightBMP cells were mixed with a MIXL1^GFP/w^ reporter cell line.^66^ Light-stimulated iRFP⁺ LightBMP cells induced neighboring MIXL1^GFP/w^ cells to adopt primitive-streak-like identity, marked by GFP expression (**Fig. 6E**). Over 2-3 days, these GFP^+^ mesodermal cells migrated away from the high-iRFP region, with some traveling hundreds of micrometers over ∼10 hours. In contrast, the region with high iRFP expression remained largely stationary, indicating that higher BMP might restrict mesodermal migration (**Fig. 6E**, **Fig. S6, Video S1**).

Together, these results demonstrate that LightBMP can spatially and temporally control BMP secretion to break symmetry within micropatterned hPSCs, generating a stripe-like domain of mesodermal differentiation. Within this domain, cells acquire mesendodermal identities, undergo EMT, and migrate anteriorly and laterally from the stimulation region, recapitulating key features of the primitive streak formation.

## DISCUSSION

Precise spatiotemporal control of morphogen signaling is essential for embryonic symmetry breaking and the establishment of developmental axes; however, current in vitro models do not fully recapitulate these processes. Here, we present LightBMP, a light-inducible platform that enables tunable, reversible, and spatially and temporally controlled expression of BMP4 (BMP) in human pluripotent stem cells (hPSCs). Using this system, we decode the rules governing BMP-driven cell-fate decisions and show that precise control of a single morphogen source is sufficient to drive symmetry breaking and model key features of primitive-streak-like patterning in a model of the epiblast disk.

Bioengineering and synthetic biology tools provide spatiotemporal control of signaling that conventional morphogen exposure cannot achieve.^67–78^ Previously reported approaches for restricting BMP signaling in space and time have key limitations: microfluidics and gradient-patterned hydrogels lack sufficient spatial and temporal precision,^79–81^ while synthetic receptor systems bypass key features of native signaling, such as ligand sensing, receptor tetramerization, and feedback regulation.^82–86^ Only a handful of studies have developed tools for the induction of extracellular ligands that preserve ligand spreading, receptor binding, and feedback regulation.^28,71,87^ A recently reported photoactivatable Cre-loxP strategy enables spatially-defined expression of extracellular BMP ligands that act on endogenous receptors, but this approach is inherently irreversible and therefore cannot probe dynamic BMP regimes.^28^ A reversible GAVPO-based system has been used to control Shh production in mouse PSCs, placing a light-induced Shh-expressing source adjacent to differentiating neural cells to generate tunable gradients and test how they are interpreted during dorsal-ventral patterning.^71^ Here, we use LightBMP to show that spatially restricted BMP expression in human pluripotent epithelium is sufficient to break symmetry and serve as the upstream trigger for gastrulation stage patterning, EMT, and directed migration.

Unlike recombinant protein treatment, LightBMP more closely mimics endogenous signaling kinetics, where pathway activation is controlled by intracellular transcriptional programs rather than exogenous ligand delivery. Continuous ligand production enables sustained, dose-dependent BMP signaling, whereas low concentrations of recombinant BMP (<3 ng/mL) generate only transient responses as the ligand is depleted from the medium, preventing prolonged low-amplitude activity.^17–19,41^ This kinetic advantage of the LightBMP platform is essential for dissecting how BMP dynamics regulate cell fate. We show that sustained low-level BMP signaling activates downstream WNT and drives mesodermal (BRA^+^) specification while remaining below the threshold for amniotic-like (ISL1^+^) differentiation, enabling mesodermal fates even after 48 hours of low-intensity illumination. At higher light intensities, LightBMP behaves as a duration-dependent morphogen, recapitulating signaling dynamics observed with higher-dose (≥3 ng/mL) recombinant BMP.^19^ Together, these findings demonstrate that the combinatorial action of BMP and the downstream WNT allows BMP to act as a morphogen across both dose and time. The key to mesoderm induction is to induce stable WNT signaling while remaining below the threshold for amnion fate, either due to low amplitude or low duration of BMP activity.

To further investigate how BMP dose influences cell-fate outcomes, we established BMP gradients across the hPSC layer by confining BMP production to one half of the culture, approximating conditions at the extraembryonic-embryonic boundary. Cells experiencing the highest BMP signaling, largely those within the source region, adopted an ISL1⁺ amnion-like identity, whereas cells below this threshold instead acquired BRA⁺ mesodermal fates. Within these BRA⁺ populations, cells progressively acquired overlapping HAND1⁺, TBX6⁺, and FOXA2⁺ identities ordered by distance from the BMP source and corresponding to lateral plate, paraxial, and axial mesoderm, respectively, mirroring the specification of these subtypes along the medial-lateral axis of the mouse embryo in response to BMP signaling.^9,11,14,52,53,88^ During gastrulation, BMP acts upstream of a morphogen cascade in which BMP induces WNT, which subsequently activates NODAL signaling.^22,89–92^ Consistently, BMP in our system induced WNT, and the WNT gradient continued to expand after the BMP gradient had stabilized, suggesting that the mesodermal domains are specified by a relayed WNT signal rather than directly by BMP. NODAL signaling was likely also activated, though we could not distinguish NODAL-mediated SMAD2/3 activation from that caused by exogenous TGF-β signaling present in the medium.^22^ Together with prior human stem cell studies, our results support a model in which higher BMP signaling promotes lateral plate mesoderm specification, moderate BMP signaling allows WNT activity to support paraxial mesoderm specification, and lower BMP/WNT signaling permits NODAL-mediated anteriorization toward axial mesodermal identities.^48,49,62,93^

In our hands, WNT activation and the emergence of primitive-streak-like cells did not require hydrogel substrates, as previously proposed.^28^ To directly test whether mechanical constraints are essential for pattern formation, we confined BMP expression within micropatterned hPSC colonies, which typically exhibit edge-driven radial differentiation.^20,21^ Under these conditions, differentiation was not dictated by colony edges; instead, lineage specification followed the spatiotemporal coordinates of BMP input. These findings align with microfluidics studies showing that basally applied BMP can drive differentiation without reliance on edge-imposed cues.^94^ Differentiation could be induced across any region of the colony, with local light dose specifying lineage identity, demonstrating that spatially restricted morphogen production can override edge-imposed patterning and is sufficient to instruct differentiation independently of mechanical constraints.

However, although mechanical constraints may not be strictly required for differentiation, they likely influence morphogen dispersal and tissue-level ligand sensing.^95^ In our system, inclusion of the ROCK inhibitor, which reduces junctional tension,^47,96^ enhanced mesodermal differentiation at the boundary between BMP-secreting and recipient cells and extended the range of pSMAD1/5/8 signaling in neighboring receiver cells. This observation is consistent with increased BMP receptor accessibility within epithelial tissues.^21,46^ However, distinguishing between the contributions of receptor accessibility and mechanosensitive signaling pathways remains challenging, as perturbations to tissue mechanics often alter both.^28,97^ Further studies will therefore be required to disentangle how mechanical properties influence morphogen interpretation and lineage patterning.

Overcoming boundary-dependent patterning addresses a key limitation of conventional micropatterned hPSC systems, in which uniform recombinant BMP treatment produces radially symmetric fate patterns that do not reflect human embryogenesis. In contrast to the concentric organization observed in vitro, gastrulation in humans and some other amniotes proceeds at one side of the epiblast disk, where the primitive streak forms as a stripe-like structure.^50,57–60^ By imposing a linear, localized low-BMP source within one side of the epiblast-like colony, we could break the radial symmetry of the disk and model a primitive-streak-like domain that more closely recapitulates human developmental patterning. This domain is defined not only by BRA⁺ and other mesodermal markers but also by morphogenetic behaviors, including epithelial-to-mesenchymal transition and directional cell migration.^2^ Notably, the emergence of primitive-streak-like patterning did not depend on colony boundary or exogenous WNT or NODAL treatments, revealing the minimal requirements for symmetry breaking by a single morphogen source.

However, we did not fully recapitulate embryonic axes organization, suggesting that additional factors come into play during development. When recreating a primitive-streak-like region within micropatterned colonies, we define the posterior at the site of primitive-streak-like formation, the anterior at the opposing side, the medial axis along the induced stripe, and lateral regions extending outward, mirroring embryonic coordinates. The posterior-anterior gradient is reproduced in our system: BMP signaling is highest at the posterior (persisting up to 20 h post-stimulation), reduced anteriorly (detectable at early time points), and largely absent at the far anterior. In contrast, the medial-lateral axis is inverted relative to embryonic conditions. In vivo, BMP is produced by surrounding extraembryonic tissues, possibly leading to increased signaling in lateral regions. In contrast, our system produces the highest BMP within the medial stripe and minimal lateral signaling, effectively inverting this axis. Future studies may address this opportunity through alternative light-patterning strategies to better reproduce embryo-like spatial organization.

Looking forward, LightBMP provides a versatile framework for probing morphogen-driven patterning across diverse developmental contexts, including cardiac specification, neural dorsoventral patterning, limb development, and angiogenesis. Its compatibility with dynamic and spatial control also makes it well-suited for integration into three-dimensional stem cell models. More broadly, this work points toward a future in which light can be used to program tissue organization with high precision in both space and time, opening new possibilities for engineering complex organoids and advancing regenerative medicine toward programmable, “on-demand” tissue and organ formation.^98^

## RESOURCE AVAILABILITY

### Lead contact

Further information and requests for resources and reagents should be directed to and will be fulfilled by the lead contact, Aryeh Warmflash.

### Materials availability

- The cell lines generated in this study are available upon request from the lead contact.
- Plasmids generated in this study have been deposited at Addgene and are publicly available as of the date of publication.

### Data and code availability

- Microscopy data reported in this paper will be shared by the lead contact upon request.
- All original code has been deposited at GitHub and is publicly available as of the date of publication.
- Any additional information required to reanalyze the data reported in this paper is available from the lead contact upon request.

## ACKNOWLEDGEMENTS

We thank Dr. Caroline Ajo-Franklin, Dr. Richard Behringer, Dr. Jeff Tabor, Dr. James Chappell, and members of the Bashor and Warmflash labs for helpful discussions. This work was supported by grants NIH R01 EB029483 (C.J.B.), NIH R35GM149328 (A.W.), R01HD112488 (A.W.), NSF MCB-2135296 (A.W.), and funding from the Robert J. Kleberg Jr and Helen C. Kleberg Foundation (C.J.B.).

## AUTHOR CONTRIBUTIONS

A.Y.S., C.J.B., and A.W. conceived of the study. A.Y.S. developed methodology, designed and performed experiments, and analyzed the data. C.J.B. and A.W. acquired funding and supervised the study. A.Y.S., C.J.B., and A.W. wrote the manuscript.

## SUPPLEMENTAL DATA FIGURE LEGENDS

**Figure S1. Design and optimization of the LightBMP circuit in hPSCs, related to Figure 1**.

**(A)** Schematic of the ePiggyBac constructs used to generate LightBMP cells. One construct drives expression of GAVPO, a blue light-inducible transcription factor composed of a Gal4 DNA-binding domain, Vivid light-sensitive dimerization domain, and p65 activation domain, with a C-terminal FLAG tag for detection. The second construct contains iRFP-P2A-proBMP4 under the control of a Gal4 upstream activation sequence (UAS) promoter. Upon blue light illumination, GAVPO dimerizes and binds UAS elements to activate transcription of iRFP-P2A-proBMP4.

**(B)** Comparison of promoter variants driving GAVPO. Fluorescence (iRFP) and immunofluorescence analysis of FLAG-tagged GAVPO and pSMAD1/5/8 in bulk transduced, antibiotic-selected hPSCs expressing iRFP-P2A-proBMP4 and GAVPO driven by TRE3G, CAG, or EF1α promoters. n = 4 images per condition.

**(C)** Immunostaining for FLAG-tagged GAVPO in bulk antibiotic-selected hPSCs harboring the TRE3G-GAVPO construct following continuous doxycycline (Dox) treatment for 10 days. Scale bar, 100 μm. n = 4 images per condition.

**(D)** Comparison of UAS promoter variants for driving iRFP-P2A-proBMP4. Basal circuit activity was assessed by measuring iRFP fluorescence in the dark (black rectangle). n = 4 images per condition.

**(E)** Clonal screening of LightBMP lines. Single-cell-derived clones (n = 21) were assessed by pSMAD1/5/8 immunostaining under light and dark conditions. Clone 10 was selected based on minimal basal activity and light responsiveness. n = 4 images per condition.

**(F)** Bright-field images of hPSCs under the indicated conditions. Wild-type cells (non-differentiated control), cells harboring a non-optimal circuit architecture (high differentiation control), and LightBMP cells cultured in the absence or presence of the BMP inhibitor NOGGIN. Representative images of the routinely cultured cells.

**(G)** Differentiation capacity toward BMP-induced ISL1⁺ fate following prolonged BMP pathway inhibition. hPSCs were treated for 10 days with LDN193189 (250 nM) or NOGGIN (125 ng/mL), followed by three PBS^−/−^ washes prior to stimulation with recombinant BMP4 (50 ng/mL) for 48 h. ISL1 expression was assessed by immunostaining. Scale bar, 100 μm. n = 4 images per condition.

**(H)** Immunostaining for pluripotency markers in the LightBMP cell line (clone 10) and parental ESI-017 hPSCs. Data represent n = 4 wells per condition, with 12 images acquired per well.

**(I)** Differentiation response of LightBMP and wild-type hPSCs to recombinant BMP. Cells were treated with indicated concentrations of BMP4 and immunostained for ISL1 (amnion) and BRA (mesoderm). Scale bar, 100 μm. n = 12 images per condition.

**Figure S2. optoBox design, calibration, and characterization of LightBMP stimulation regimes, related to Figure 2**.

**(A)** Cross-section schematic of the optoBox for a single well. A visible-light bandpass filter (400–700 nm) is placed ∼4 mm above each blue LED (465 nm peak) to reduce infrared heat transfer while preserving blue-light transmission.

**(B)** Temperature increase of cell culture medium under continuous illumination. Measurements were performed with and without a heat sink or optical filter. The data represent n = 6 measurements randomly sampled across wells of the 96-well plate.

**(C)** Calibration of optoBox light output. Arduino-programmed intensity units (bytes) are linearly proportional to measured irradiance. Light intensity was measured at the bottom of empty wells across a range of programmed inputs with the full optoBox assembly (including optical filters). Data represent n = 3 wells per intensity condition.

**(D)** Cell viability as a function of light input. Percentage of PI (propidium iodide)–positive cells following 24 h of continuous illumination, plotted against total light dose (bottom x-axis) and corresponding light intensity (top x-axis). n = 3 images per condition.

**(E)** Time course of BMP signaling under pulsed illumination. pSMAD1/5/8 immunofluorescence and iRFP expression were quantified during pulsed light stimulation (10 s pulses every 100 s; ON) and following cessation of illumination after 24 h (OFF) at the indicated light intensities. n = 5 images per condition.

**(F)** Relationship between activated and responding cells. Fraction of light-activated (iRFP⁺) cells versus responding (pSMAD1/5/8⁺) cells, related to Figure 2C. n = 4 images per condition.

**Figure S3. Light-induced low BMP signaling promotes sustained mesodermal differentiation, related to Figure 3**.

**(A)** Percentage of BRA⁺ cells across light intensities, related to Figure 3B. Cells were stimulated under pulsed illumination (10 s every 100 s) in the presence of ROCK inhibitor (10 µM) and fixed at 48 h. n = 4 images per condition.

**(B)** Percentage of BRA⁺ cells across light intensities in the absence of ROCK inhibitor. Cells were stimulated under pulsed illumination (10 s every 100 s) and fixed at 48 h. n = 4 images per condition.

**(C)** Analysis of WNT and BMP pathway activity across stimulation regimes. Cells were stimulated for 21, 32, or 48 h at the indicated light intensities and immunostained for LEF1 (WNT activity), pSMAD1/5/8 (BMP signaling), and lineage markers. At each duration, both mesodermal (BRA⁺) and amnion-like (ISL1⁺) conditions exhibit high WNT signaling, whereas higher BMP signaling levels correspond to ISL1 induction. pSMAD1/5/8 levels should not be compared across columns due to signal decay after illumination (indicated by the green arrow at the bottom). n = 4 images per condition.

**(D)** Immunostaining of additional pluripotency and lineage markers across LightBMP stimulation regimes. Cells were stimulated for varying durations (time) and light intensities (dose), then fixed at 48 h. Shown are NANOG (pluripotency), MIXL1 (mesoderm), and GATA3 (amnion), related to Figure 3C. Scale bar, 100 μm. n = 16 images per condition.

**Figure S4. Spatial BMP gradients and mesodermal patterning in half-illuminated LightBMP cultures, related to Figure 4**.

**(A)** iRFP and DAPI channels for the indicated conditions in Figures 4 and S4. The white dotted line denotes the boundary defined by iRFP⁺ cells (computationally segmented). Scale bar, 1000 μm.

**(B)** Effect of ROCK inhibition on BMP signaling range and mesodermal differentiation. Comparison of pSMAD1/5/8, LEF1, BRA, and ISL1 immunostaining in half-illuminated cultures with or without ROCK inhibitor during 2 days of illumination. Scale bar, 100 μm. n = 3 image grids per condition.

**(C)** Spatial organization of mesodermal transcription factors along the BMP gradient after 3 days of light stimulation. Immunostaining for HAND1 (lateral mesoderm), TBX6 (paraxial mesoderm), and FOXA2 (axial mesoderm). Scale bar, 1000 μm. Data represent n = 3 (1 × 3 image grids) per condition.

**(D)** Immunostaining of additional lineage markers after 4 days of half-well illumination. Scale bar, 1000 μm. n = 3 (1 × 3 image grids) per condition across 2 wells.

**Figure S5. Light-patterned BMP secretion generates gastrulation-like organization in the absence of colony edge constraint, related to Figure 5**.

**(A)** Generation of ring-shaped BMP secretion using laser-scanning (488 nm, 10 s per ring every 450 s) illumination. iRFP expression and pSMAD1/5/8 immunostaining after 2 days of ring-patterned stimulation at the indicated light intensities. Scale bar, 500 μm. n = 3 ring patterns per condition.

**(B)** Fate patterning around light-induced BMP rings. Immunostaining for ISL1, BRA, and SOX2 at day 3 following 2 days of stimulation under the indicated illumination intensities (488 nm, 10 s per ring every 450 s). Scale bar, 500 μm. Data represent n = 3 ring patterns per condition.

**Figure S6. Migration of a representative primitive streak-like cell, related to Figure 6E**.

Sparse MIXL1^GFP/w^ reporter cells (yellow), mixed with LightBMP cells, are induced near the iRFP⁺ stripe (light-activated BMP secretors) and migrate away over time. A representative single-cell migration track is shown; the orange arrow (manually annotated) indicates the tracked cell, while migration tracks were computationally derived. Scale bar, 100 μm.

## STAR★METHODS

### EXPERIMENTAL MODEL AND STUDY PARTICIPANT DETAILS

#### Cell lines

The ESI-017 human pluripotent stem cell (hPSC) line (RRID: CVCL_B854) was obtained from ESIBIO, with karyotype and genomic integrity validated by the manufacturer. The LightBMP cell line was generated from the parental ESI-017 line. For cell-tracking experiments, the MEL1-MIXL1^GFP/w^ reporter line,^66^ a gift from Ed Stanley (Murdoch Children’s Research Institute), and a LightBMP-mCherry-CAAX line (generated from the parental LightBMP) were used. All cell lines were routinely tested for mycoplasma contamination and confirmed negative. All experiments were conducted in accordance with the ISSCR guidelines found at https://www.isscr.org/policy/guidelines-for-stem-cell-research-and-clinical-translation.

### METHOD DETAILS

#### Cell Culture

Cells were maintained in mTeSR Plus medium (STEMCELL Technologies) on tissue culture dishes coated with Matrigel (Corning) or Geltrex (Life Technologies), using a 1:200 dilution in DMEM/F12 (Corning). Coatings were prepared either overnight at 4 °C or for 1 h at 37 °C. LightBMP cells were supplemented with 125 ng/mL NOGGIN (R&D Systems) to prevent differentiation during extended culture. Cells were passaged every 3-4 days using Dispase (STEMCELL Technologies). For experiments, cells were harvested at 70–80% confluence and dissociated into a single-cell suspension using Accutase (Corning). After plating, the ROCK inhibitor Y-27632 (10 µM; STEMCELL Technologies) was added to support cell viability.

#### DNA assembly

Plasmids were assembled using a custom hierarchical Golden Gate cloning strategy, as previously reported.^99,100^ Briefly, DNA fragments and oligonucleotides were commercially synthesized (Twist Bioscience or IDT) and cloned into carbenicillin-resistant entry vectors using BsaI-HFv2 (NEB) to generate ‘Level 1’ part plasmids (promoters, open reading frames [ORFs], and terminators). Promoter parts included constitutive pCAG and pEF1α (Table S1), the doxycycline-inducible TRE3G promoter (Table S3), and Gal4 upstream activating sequences (UAS)-based promoters 5×UAS-TATA, 5×UAS-AMLP, and 3×UAS-AMLP (Table S2). ORF assemblies included GAVPO and iRFP-P2A-proBMP. The GAVPO sequence was obtained from Wang et al. (2012),^29^ iRFP670 from Shcherbakova and Verkhusha (2013),^30^ and proBMP4 from the NCBI GenBank (Accession No. NM_130850). Amino acid sequences for all encoded proteins are provided in Table S4. The SV40 polyadenylation signal was used as the terminator. ‘Level 2’ expression units were assembled by combining part plasmids into kanamycin-resistant vectors using Esp3I (Thermo Fisher Scientific). ‘Level 3’ assemblies were generated using BbsI-HF (NEB) by cloning expression units into ePiggyBac destination vectors: ePB-UbC-rtTA-T2A-PuroR for all GAVPO constructs and ePB-UbC-BlastR-FloxP-EBFP2 for all iRFP-P2A-proBMP constructs. All plasmids were verified by restriction digest and whole-plasmid sequencing (Oxford Nanopore Technologies).

#### DNA integration

Transfected plasmids were prepared using a HiSpeed Plasmid Midi Kit (Qiagen) and diluted to 1 µg/mL (0.65 µg/mL for the helper transposase plasmid) based on NanoDrop (Thermo Fisher Scientific) measurements. DNA was delivered by electroporation using the Neon Transfection System (Invitrogen, 10 µL kit). For each reaction, 2 × 10⁵ cells were resuspended in 9.5 µL Buffer R and mixed with 1 µL DNA containing ePiggyBac helper plasmid (0.2 µL), along with ePB-GAVPO (0.4 µL), and ePB-iRFP-P2A-proBMP4 (0.4 µL) expression constructs. Electroporation was performed at 1200 V, 10 ms, with 3 pulses, and cells were immediately seeded onto pre-coated 24-well plates. Antibiotic selection was initiated 3 days post-transfection using puromycin (5 µg/mL) and blasticidin (5 µg/mL) and maintained for 2 weeks with daily medium changes.

#### Generation of monoclonal cell lines

The antibiotic-enriched bulk population carrying TRE3G-GAVPO-FLAG and 3×UAS_AMLP-iRFP-P2A-proBMP transgenes was induced with doxycycline (1 µg/mL) and light for 18 h in the presence of LDN193189 (250 nM) (Thermo Fisher Scientific). iRFP-positive cells were single-cell sorted into 96-well plates using a SH800S Cell Sorter (Sony) equipped with Cell Sorter Software (version 2.2.4.5150). Cells were cultured in mTeSR Plus supplemented with 1× CloneR, with LDN193189 (250 nM) maintained during induction and for 48 h post-sorting to prevent differentiation. Following expansion, 21 surviving undifferentiated clones were evaluated by pSMAD1/5/8 immunostaining upon constant light and dark conditions (**Figure S1E**). A light-responsive LightBMP clone with minimal basal signaling (clone 10) was selected for further experiments. Experiments for Figs. 2B-D, 3B, 6C were repeated with an independent clone (clone 7), yielding consistent results.

#### optoBox construction

For high-throughput optogenetic stimulation (**Figs. 1-4**), blue light (peak emission 465 nm) was delivered using a custom-built device (“optoBox”) incorporating a 96-LED matrix (MapleBear Electronics)^37^. The optoBox assembly included a 96-well adapter,^38^ customized to fit the optoBox (CAD file available on GitHub), and mounted to the LED matrix using heat-set insertion. To ensure thermal management, an aluminum heat sink (120 × 69 × 27 mm; Awxlumv) was attached to the underside of the LED board using thermal paste (Arctic MX-5). Two cooling fans (60 × 10 mm; StarTech, FAN6X1TX3) were mounted to the heat sink fins to dissipate heat, and the assembly was elevated on a custom stage to allow airflow beneath the device. Visible bandpass filters (IR-cut, 8 × 8 × 0.7 mm; Quanmin) were positioned within each well of the adapter using integrated supports that held the filters ∼4 mm above each LED, reducing heat transfer to the cell culture (**Fig. S2A-B**). The LED board and microcontroller (D1 Mini, Wemos Electronics) were powered via a 5 V supply through a capacitor- and resistor-based power board (Maple Electronics). Cooling fans were powered at 12 V.

LED control programs (Arduino IDE) were validated using short-duration test regimes and included an EEPROM-based recovery loop to restore timing if microcontroller resets during multiday experiments. Light intensity (power density) was measured at the bottom of wells using a power meter (PM20A; Thorlabs) with a fiber probe (M59L01; 480 nm; n = 3), confirming a linear relationship between programmed input and delivered intensity (**Figure S2C**).

#### High-throughput optogenetic stimulation

Optogenetic experiments (**Figs. 1-4**) were performed in black-walled 96-well plates (Corning) with 2 × 10⁴ cells seeded in 150 µL of mTeSR Plus medium (STEMCELL Technologies) plus doxycycline (1 μg/mL, Sigma-Aldrich) and ROCK inhibitor Y-27632 (10 µM; STEMCELL Technologies) in each well 24 h prior to stimulation. If not stated differently, doxycycline and ROCK inhibitor were retained for the duration of the experiment.

For ON-kinetics experiments (**Figs. 2B, S2E**), illumination was programmed such that all samples were fixed simultaneously (e.g., an 8 h condition received 8 h of illumination immediately prior to fixation).

For gradient formation experiments, light leakage into dark regions was minimized by applying light-blocking adhesive tape (TygoTy) to the plate bottom, leaving only half of each well exposed to light. The tape was removed prior to imaging. ROCK inhibitor was removed after the first 48 h of stimulation.

#### Laser-scanning patterned stimulation

Low-throughput optogenetic experiments (**Figs. 5B, S5A–S5B**) were performed in glass-bottom, black-walled 96-well plates (Ibidi), with 3 × 10⁴ cells seeded in 150 µL per well 24 h prior to stimulation in the presence of doxycycline (1 µg/mL, Sigma-Aldrich). Experiments in **Figs. 5C** and **6** were conducted in 2D micropatterned cultures using 96-well black-walled, glass-bottom microplates (CYTOO). ROCK inhibitor Y-27632 (10 µM; STEMCELL Technologies) was retained for the time of stimulation. Optogenetic stimulation was delivered using an Olympus FV3000 laser-scanning confocal microscope equipped with a 20× objective (NA 0.75), a 488 nm laser, and FV31S-SW software (v2.4.1.198, Evident Scientific). To minimize light scattering and off-target activation, 50% light-dimming adhesive tape (Jieheng) was applied to the bottom of wells during stimulation and removed prior to imaging. Light intensity was measured at the bottom of taped wells (488 nm) using a power meter (PM20A with M59L01, Thorlabs) and varied linearly with laser power (intensity = 2.4562 × laser power). The following moving point laser regimes were used: **Fig. 5B**, 10-s scan every 150 s for 48 h at varying intensities; **Fig. 5C**, 10-s scan every 300 s for 48 h at varying intensities; **Fig. S5**, 10-s scan every 450 s for 48 h at varying intensities; **Fig. 6B-E** and **S6**, 1.5 μW/mm² light delivered in a spatially restricted stripe, with a 5-s scan per stripe segment repeated every ∼200 s for ∼24 h.

#### 2D micropatterned culture

Micropattern experiments were performed in 96-well glass-bottom microplates (CYTOO) with circular micropatterns 700 μm in diameter, according to the protocol described by Deglincerti et al. (2016).^101^ Wells were pre-coated with 200 µL of 5 µg/mL LN-521 (Biolamina; diluted in PBS+/+) and incubated at 37 °C for 2 h. Wells were then washed with PBS+/+ (Cytiva) using nine partial washes (150 µL removal/addition) followed by one complete wash. Cells were dissociated into a single-cell suspension using Accutase and seeded at 1× 10^5^ cells per well in 200 µL mTeSR Plus supplemented with ROCK inhibitor (10 µM) and NOGGIN (250 ng/mL). Plates were incubated at 37 °C for 40 min to allow cell attachment. Unattached cells were removed by gentle multiple washes with PBS−/− (Cytiva). After washing, 200 µL of mTeSR Plus containing ROCK inhibitor (10 µM), NOGGIN (250 ng/mL), and doxycycline (1 µg/mL) was added to each well. Cells were incubated for 24 h, washed twice with PBS−/−, and the medium was replaced with mTeSR Plus containing doxycycline (1 µg/mL).

#### Differentiation with recombinant BMP4

At 24 h after seeding, cells were treated with defined concentrations of recombinant BMP4 (R&D Systems) in mTeSR Plus medium containing ROCK inhibitor (10 µM) for 48 h in the absence of doxycycline and under dark conditions. For micropatterned BMP control experiments, ROCK inhibitor was withdrawn, and BMP4 was added at a final concentration of 50 ng/mL for 48 h.

#### Cell death assay

Cells were incubated with Hoechst 33342 (20 µM final concentration; 1:1000, Thermo Fisher Scientific) and propidium iodide (5 µg/mL, Thermo Fisher Scientific) added directly to the culture medium for 15 min at 37 °C. Plates were then centrifuged for 2 min at 162 × g (applied from both orientations) to sediment non-adherent dead cells. Imaging was performed using continuous ZDC autofocus for rapid acquisition.

#### Immunofluorescence staining

Cells were fixed in 4% paraformaldehyde (Electron Microscopy Sciences; prepared in PBS-/-) for 20 min at room temperature, followed by two washes with PBS-/-. Samples were then incubated in blocking solution (PBS-/- containing 0.1% Triton X-100 and 3% donkey serum) for 1 h at room temperature. Primary antibodies were diluted in blocking solution as listed in Table S5 and incubated with samples overnight at 4 °C. Samples were subsequently washed four times for 5 min each with PBST (PBS-/- supplemented with 0.1% Tween 20). Secondary antibodies (dilutions listed in Table S5) and DAPI (1:5000) were then applied for 1 h at room temperature. Following four additional 5 min washes with PBST, samples were maintained in PBS-/- until imaging.

#### Fixed-cell imaging

Imaging was performed using an Olympus FV3000 equipped with FV31S-SW. Images were acquired with 10× (NA 0.40) or 20× (NA 0.75) objectives. For half-well illumination experiments, imaging positions were predefined using continuous ZDC prior to the removal of the light-blocking adhesive tape. Positions were selected to capture the illuminated region and adjacent dark regions in a 1 × 3 grid; three such grids were defined per well, and the wells were imaged after tape removal. For standard cultures, at least four fields of view were acquired per well, with replication across wells where possible. For micropatterned cultures, at least four colonies were imaged per condition.

#### Live-cell imaging and single-cell tracking

CYTOO micropatterned 96-well plates were imaged using an Olympus FV3000 laser scanning confocal microscope equipped with an environmental chamber maintained at 37 °C, ∼50% humidity, and 5% CO₂. Images were acquired using a 20× air objective (NA 0.75). Cell tracking was performed in Ilastik^102^ using the Tracking with Learning workflow, applied independently to the MIXL1 and iRFP channels. Resulting tracks were further analyzed using a custom Python script.

### QUANTIFICATION AND STATISTICAL ANALYSIS

#### Image analysis

Maximum-intensity projections and image stitches were generated using FV31S-SW software. Nuclear segmentation was performed in Ilastik (version 1.4.0)^102^ using the Pixel Classification workflow to generate binary nuclear masks in the DAPI channel; in experiments where SOX2 was imaged in the 405 nm channel, a combination of nuclear markers was used to define nuclear regions. Subsequently, the Object Classification workflow was used across all channels to extract mean nuclear intensities and, where applicable, cytoplasmic intensities from pixels surrounding the nuclear masks. Resulting data were exported as dataframes and analyzed using custom scripts in MATLAB (R2024b, MathWorks) or Python 3.10 / JupyterLab v4.4.4. For micropatterned colony analyses, only nuclear masks were used for downstream processing in Python. When available, fluorescence intensities were normalized to the DAPI channel to account for variations in cell density. For visualization, look-up tables were adjusted in Fiji,^103^ and, where necessary, outlier removal filters were applied to raw images to reduce non-specific bright puncta.

#### Statistics and reproducibility

For each image, mean nuclear intensity was calculated by averaging of single-cell nuclear intensities (mean across cells). For each condition, values are reported as mean ± SD across n images. At least three images were acquired per condition. Experiments were repeated three times and yielded consistent results.

### KEY RESOURCES TABLE

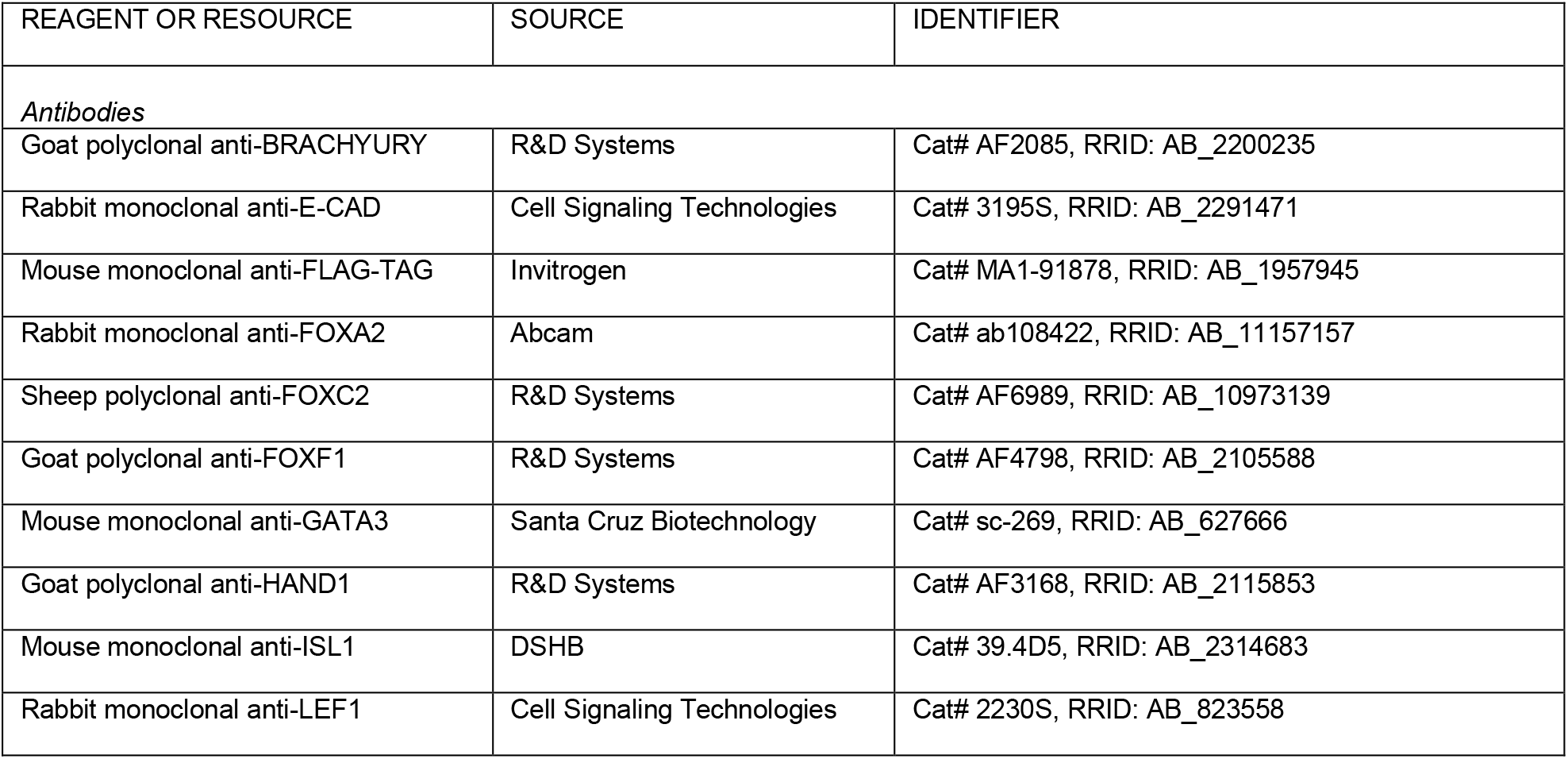
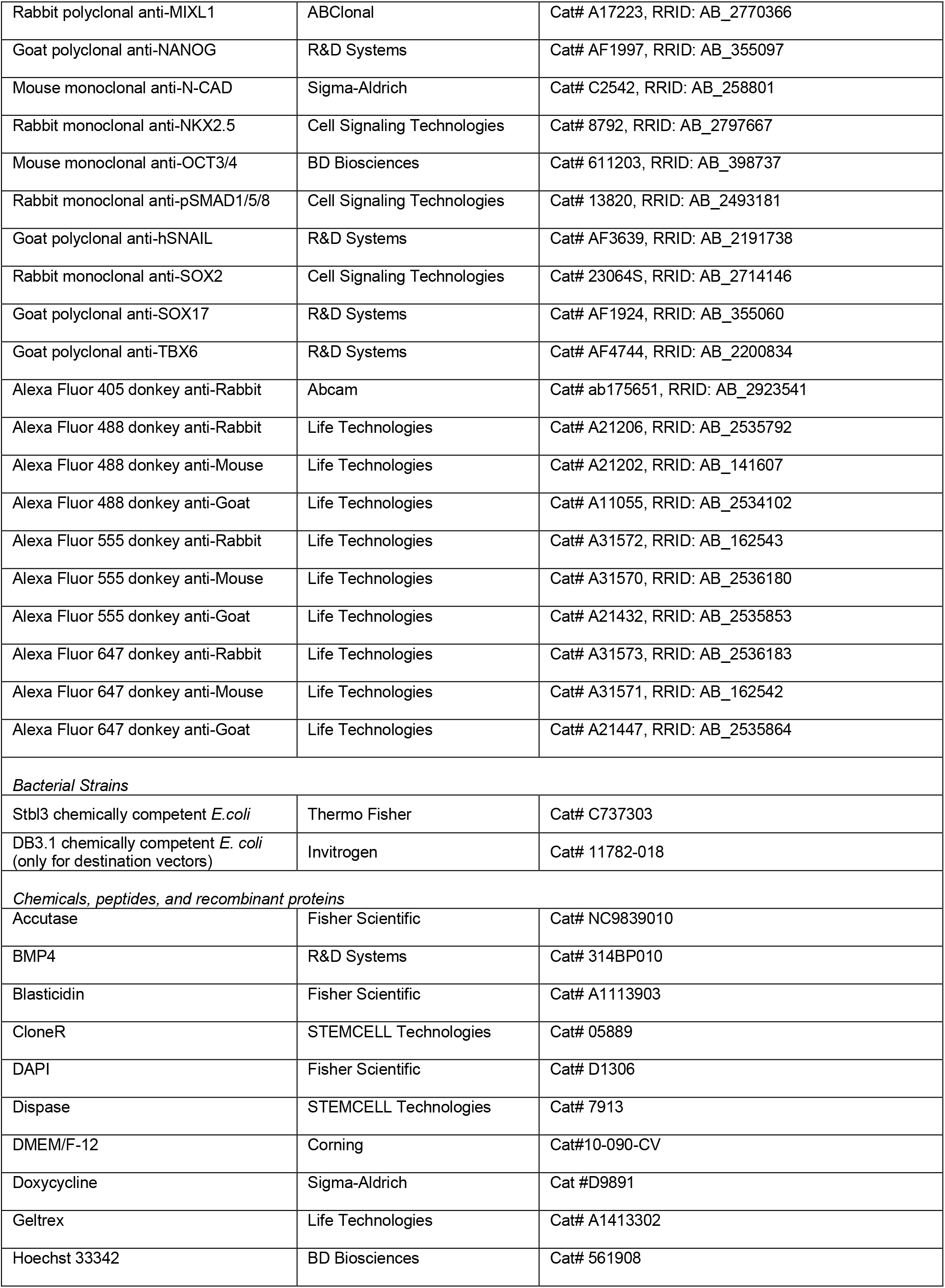
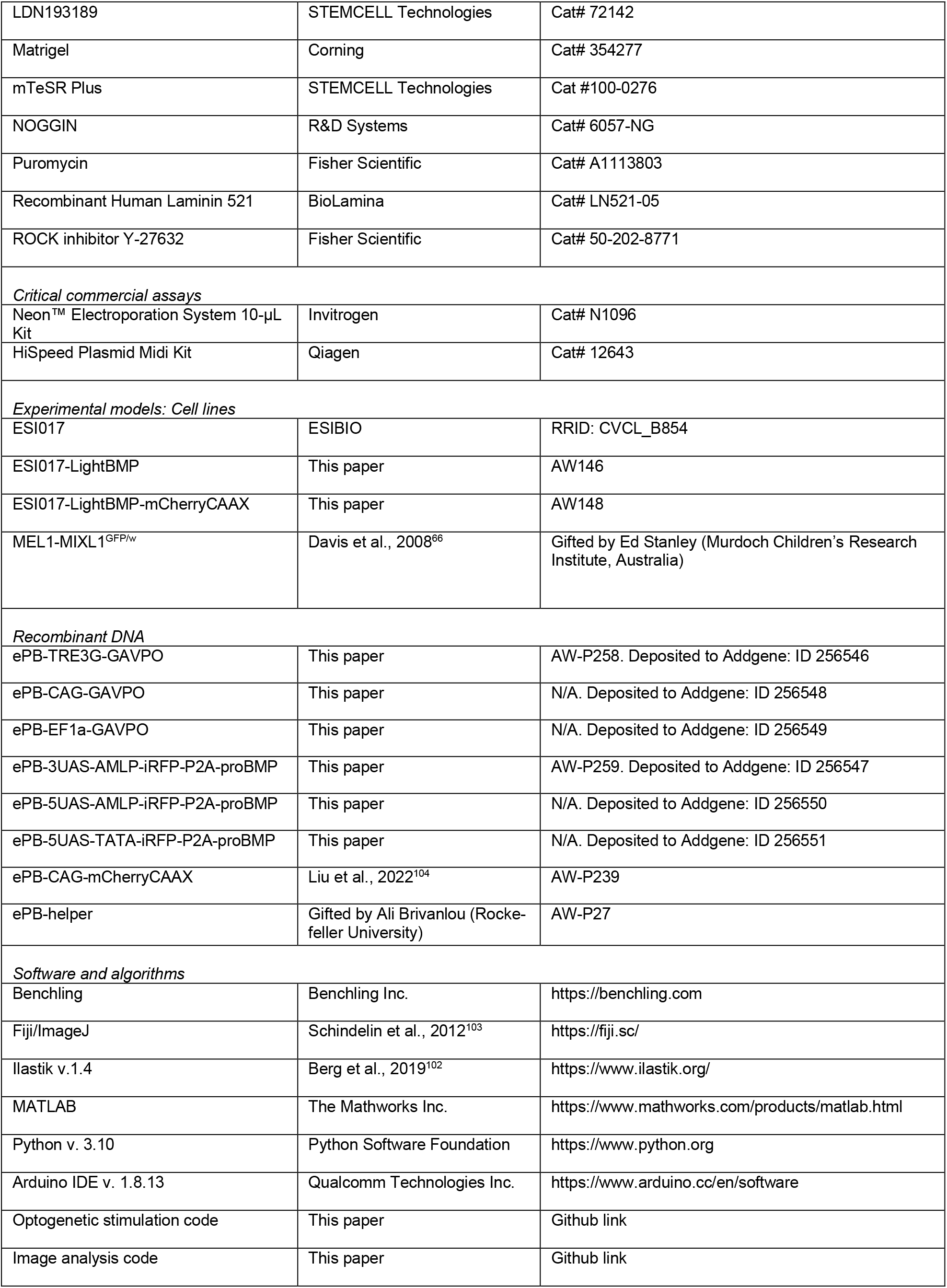

### SUPPLEMENTARY INFORMATION

**Document S1. Tables S1–S5 and supplemental references**

**Video S1. Primitive-streak-like cell migration, related to Figure 6**

MIXL1^GFP/w^ cells (yellow) migrate from the linear light-induced region (iRFP^+^ cells). Scale bar, 100 μm

