## Supplementary material for "Optogenetic BMP4 models symmetry breaking and primitive streak formation in hPSCs": Figure S1-S6

**Fig S1. Design and optimization of the LightBMP circuit in hPSCs, related to Figure 1.**

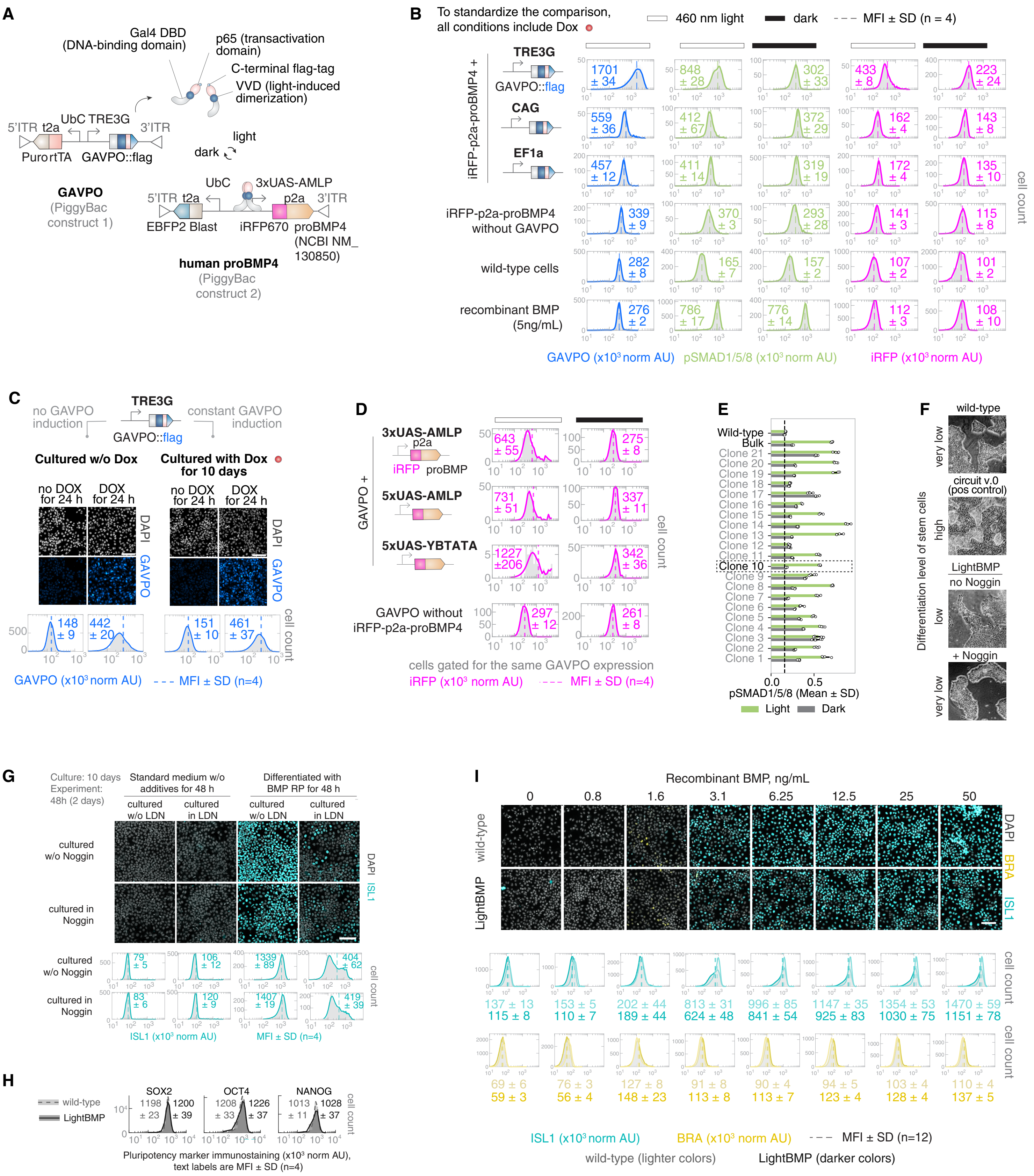

**Fig S2. optoBox design, calibration, and characterization of LightBMP stimulation regimes, related to Figure 2.**

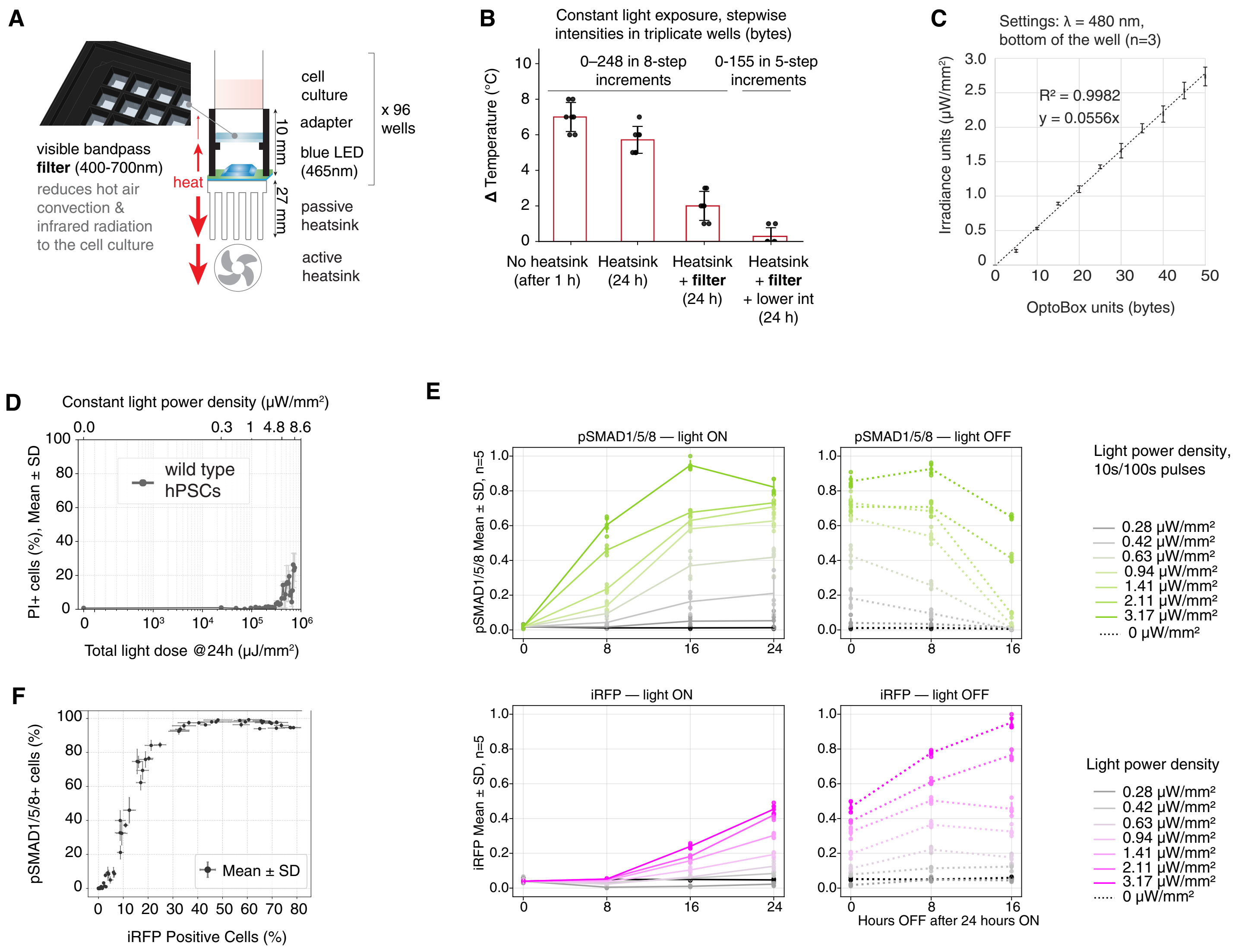

**Fig S3. Light-induced low BMP signaling promotes sustained mesodermal differentiation, related to Figure 3.**

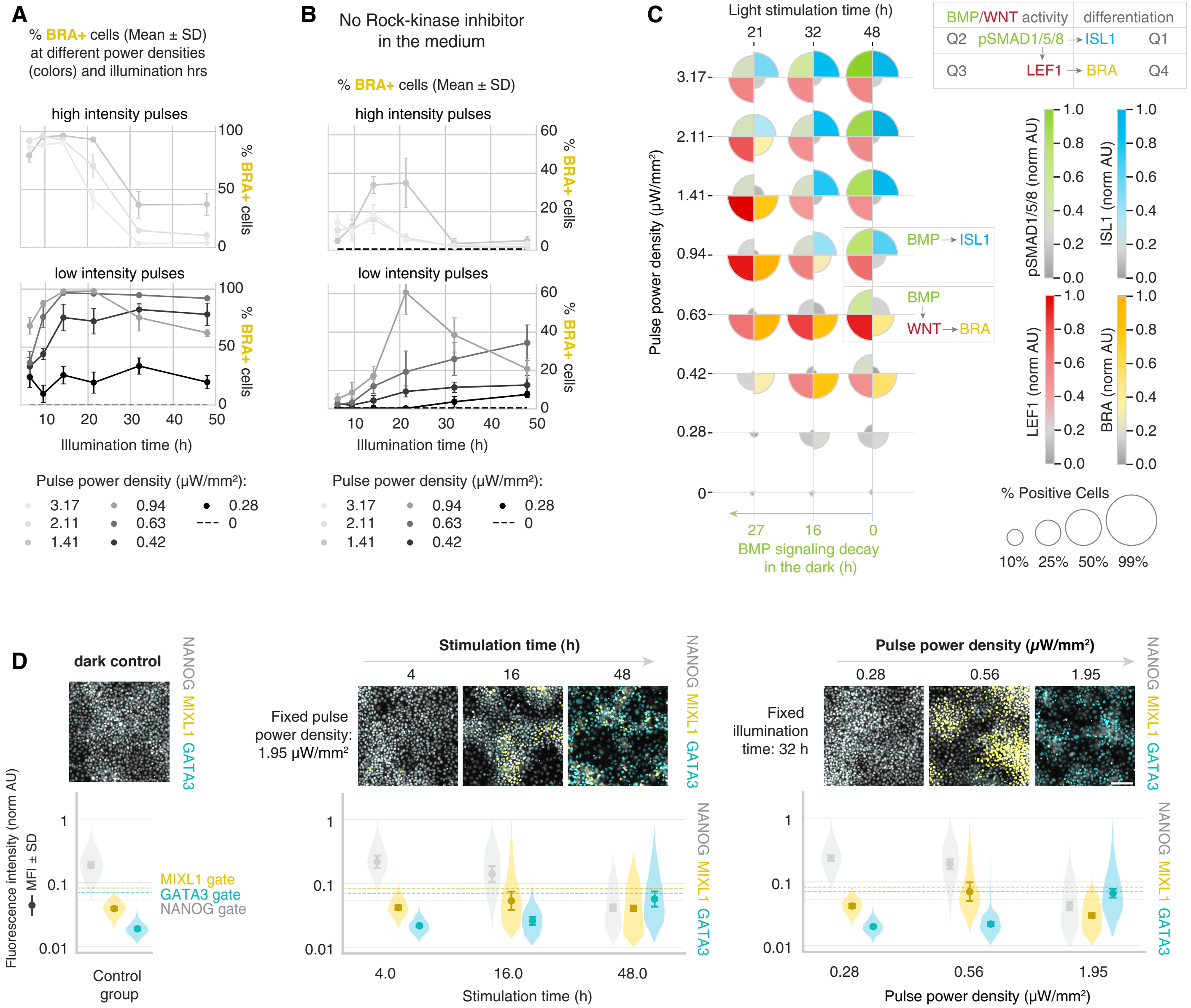

### Fig S4. Spatial BMP gradients and mesodermal patterning in half-illuminated LightBMP cultures, related to Figure 4.

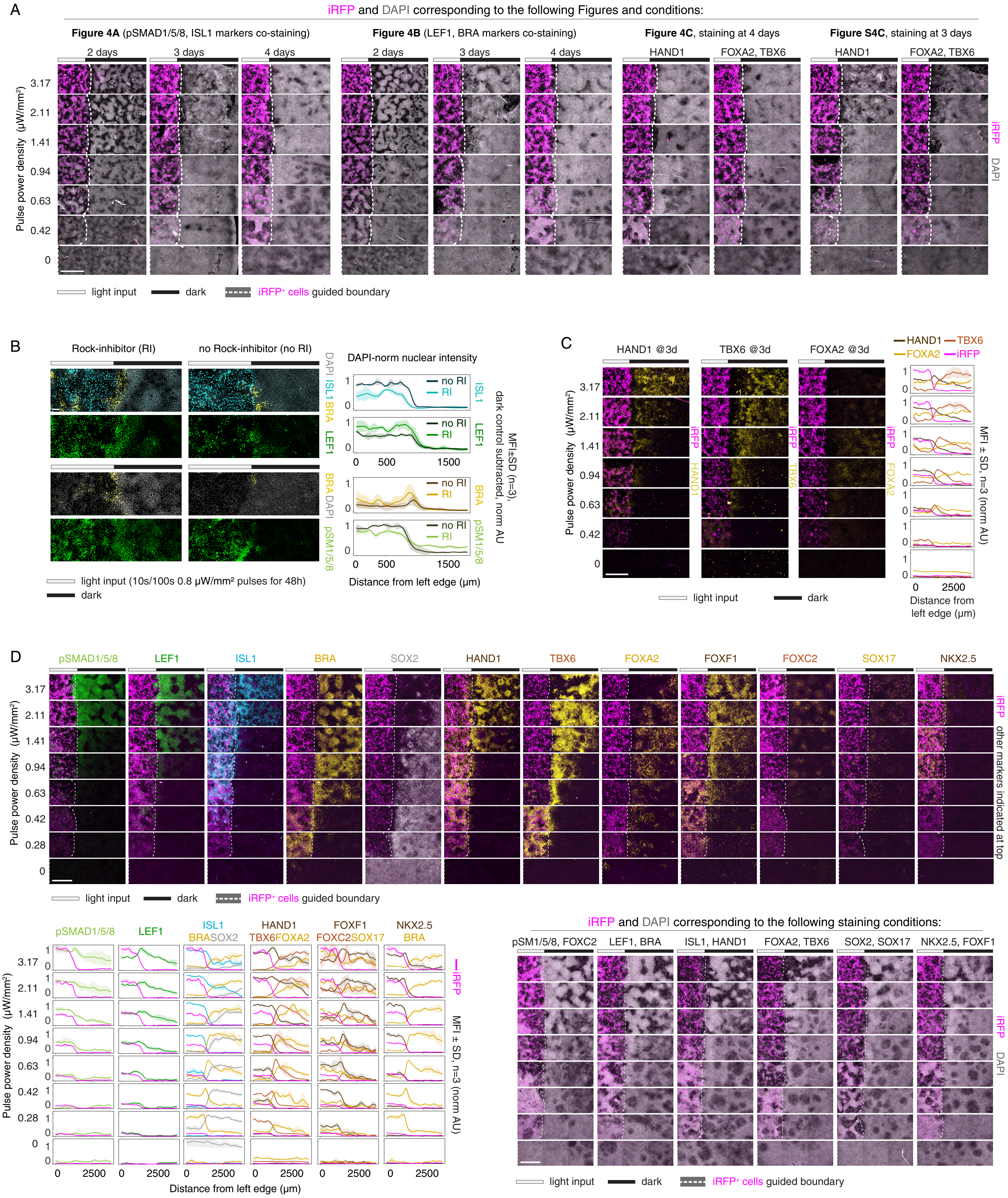

**Fig S5. Light-patterned BMP secretion generates gastrulation-like organization in the absence of colony edge constraint, related to Figure 5.**

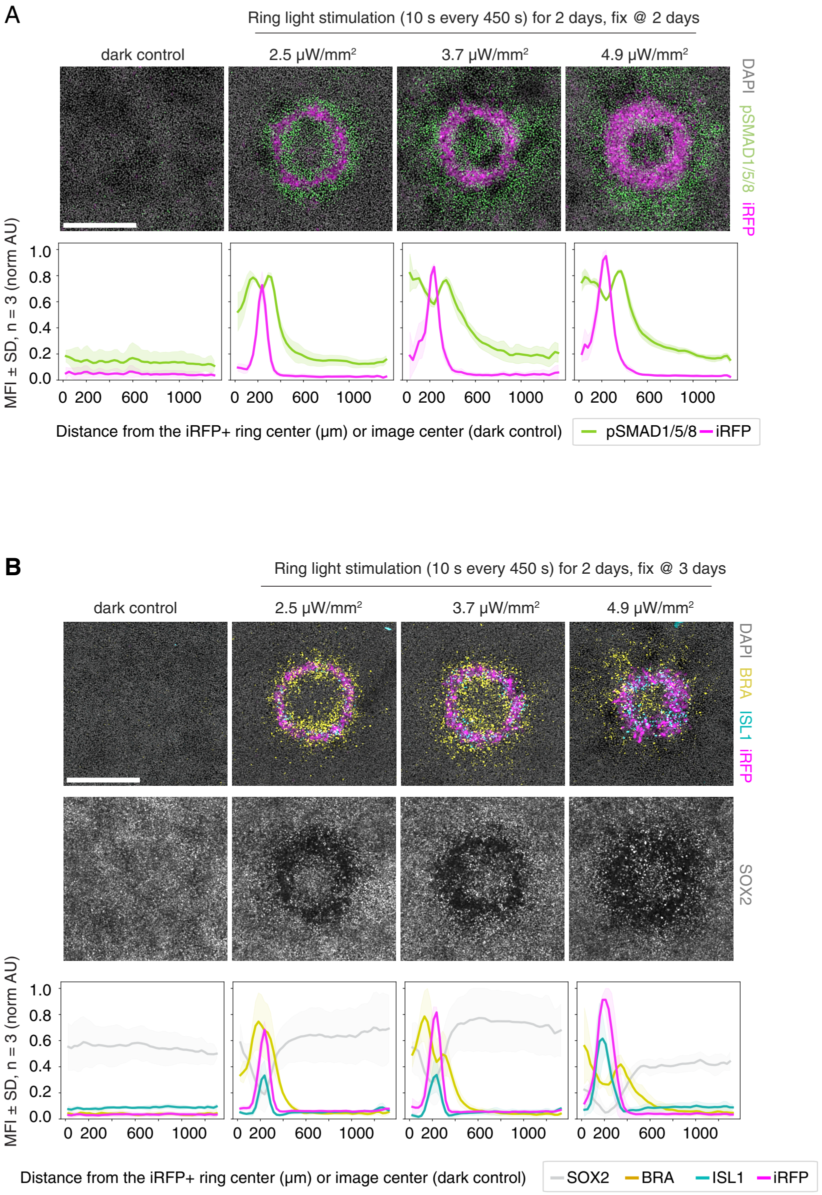

### Fig S6. Migration of a representative primitive streak-like cell, related to Figure 6E.

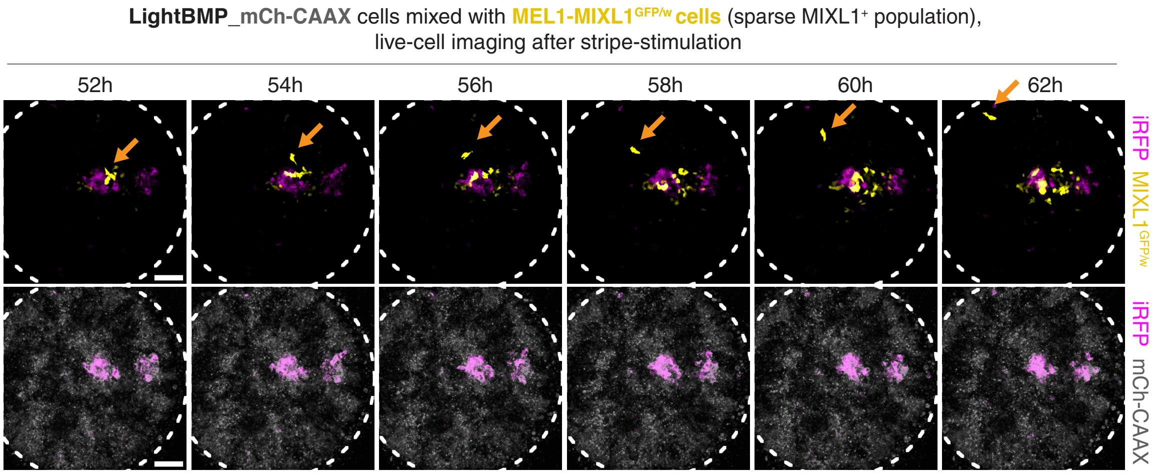


Scale bar, 100 µm

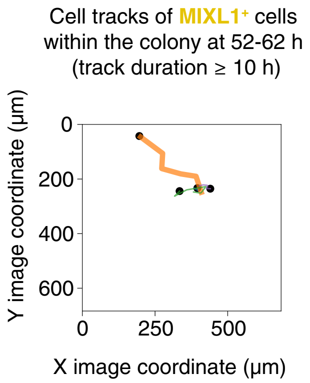
