## Supplementary material for "Optogenetic BMP4 models symmetry breaking and primitive streak formation in hPSCs": Tables S1-S5

| Promoter | Promoter Sequence |
|---|---|
| CAG | CTCGACATTGATTATTGACTAGTTATTAATAGTAATCAATTACGGGGTCATTAGTTCATAGCCCATATATGG<br>AGTTCCGCGTTACATAACTTACGGTAAATGGCCCGCCTGGCTGACCGCCCAACGACCCCCGCCCATTGA<br>CGTCAATAATGACGTATGTTCCCATAGTAACGCCAATAGGGACTTTCCATTGACGTCAATGGGTGGAGTA<br>TTTACGGTAAACTGCCCACTTGGCAGTACATCAAGTGTATCATATGCCAAGTACGCCCCCTATTGACGTC<br>AATGACGGTAAATGGCCCGCCTGGCATTATGCCCAGTACATGACCTTATGGGACTTTCCTACTTGGCAGT<br>ACATCTACGTATTAGTCATCGCTATTACCATGGTCGAGGTGAGCCCCACGTTCTGCTTCACTCTCCCCATC<br>TCCCCCCCCTCCCCACCCCCAATTTTGTATTTATTTATTTTTTAATTATTTTGTGCAGCGATGGGGGCGGG<br>GGGGGGGGGGGGCGCGCGCCAGGCGGGGCGGGGCGGGGCGAGGGGCGGGGCGGGGCGAGGC<br>GGAGAGGTGCGGCGGCAGCCAATCAGAGCGGCGCGCTCCGAAAGTTTCCTTTTATGGCGAGGCGGC<br>GGCGGCGGCGGCCCTATAAAAAGCGAAGCGCGCGGCGGGCGGGAGTCGCTGCGCGCTGCCTTCGCC<br>CCGTGCCCCGCTCCGCCGCCGCCTCGCGCCGCCCGCCCCGGCTCTGACTGACCGCGTTACTCCCACAG<br>GTGAGCGGGCGGGACGGCCCTTCTCCTCCGGGCTGTAATTAGCGCTTGGTTTAATGACGGCTTGTTTCT<br>TTTCTGTGGCTGCGTGAAAGCCTTGAGGGGCTCCGGGAGGGCCCTTTGTGCGGGGGGAGCGGCTCG<br>GGGGGTGCGTGCGTGTGTGTGTGCGTGGGGAGCGCCGCGTGCGGCTCCGCGCTGCCCGGCGGCTGT<br>GAGCGCTGCGGGCGCGGCGCGGGGCTTTGTGCGCTCCGCAGTGTGCGCGAGGGGAGCGCGGCCGG<br>GGGCGGTGCCCCGCGGTGCGGGGGGGGCTGCGAGGGGAACAAAGGCTGCGTGCGGGGTGTGTGCG<br>TGGGGGGGTGAGCAGGGGGTGTGGGCGCGTCGGTCGGGCTGCAACCCCCCCTGCACCCCCCTCCCC<br>GAGTTGCTGAGCACGGCCCGGCTTCGGGTGCGGGGCTCCGTACGGGGCGTGGCGCGGGGCTCGCCG<br>TGCCGGGCGGGGGGTGGCGGCAGGTGGGGGTGCCGGGCGGGGCGGGGCCGCCTCGGGCCGGGGA<br>GGGCTCGGGGGAGGGGCGCGGCGGCCCCCGGAGCGCCGGCGGCTGTCGAGGCGCGGCGAGCCGC<br>AGCCATTGCCTTTTATGGTAATCGTGCGAGAGGGCGCAGGGACTTCCTTTGTCCCAAATCTGTGCGGAG<br>CCGAAATCTGGGAGGCGCCGCCGCACCCCCTCTAGCGGGCGCGGGGCGAAGCGGTGCGGCGCCGGC<br>AGGAAGGAAATGGGCGGGGAGGGCCTTCGTGCGTCGCCGCGCCGCCGTCCCCTTCTCCCTCTCCAGC<br>CTCGGGGCTGTCCGCGGGGGGACGGCTGCCTTCGGGGGGGACGGGGCAGGGCGGGGTTCGGCTTC<br>TGGCGTGTGACCGGCGGCTCTAGAGCCTCTGCTAACCATGTTCATGCCTTCTTCTTTTTCCTACAGCTCC<br>TGGGCAACGTGCTGGTTATTGTGCTGTCTCATCATTTTGGCAAAGAATTGCAAGTTTGTACAAAAAAGC<br>AGGCTGAAT |
| EF1α | GGATCTGCGATCGCTCCGGTGCCCGTCAGTGGGCAGAGCGCACATCGCCCACAGTCCCCGAGAAGTTG<br>GGGGGAGGGGTCGGCAATTGAACGGGTGCCTAGAGAAGGTGGCGCGGGGTAAACTGGGAAAGTGA<br>TGTCGTGTACTGGCTCCGCCTTTTTCCCGAGGGTGGGGGAGAACCGTATATAAGTGCAGTAGTCGCCGT<br>GAACGTTCTTTTTCGCAACGGGTTTGCCGCCAGAACACAGCTGAAGCTTCGAGGGGCTCGCATCTCTC<br>CTTCACGCGCCCGCCGCCCTACCTGAGGCCGCCATCCACGCCGGTTGAGTCGCGTTCTGCCGCCTCCCG<br>CCTGTGGTGCCTCCTGAACTGCGTCCGCCGTCTAGGTAAGTTTAAAGCTCAGGTCGAGTCCGGGCCTTT<br>GTCCGGCGCTCCCTTGGAGCCTACCTAGACTCAGCCGGCTCTCCACGCTTTGCCTGACCCTGCTTGCTC<br>AACTCTACGTCTTTGTTTCGTTTTCTGTTCTGCGCCGTTACAGATCCAAGCTGTGACCGGCGCCTACGAA<br>T |

**Table S1.** Constitutive promoters tested for GAVPO expression (related to **Fig. S1B**). The sequence of the doxycycline-inducible promoter is provided in Table S3.

| Part Class | Part Name | Part Sequence |
|---|---|---|
| Enhancer sites | n = 3 x UAS binding sites | ATCGATAGGTACCGAGTTTCTAGACGGAGTACTGTCCTCCGAGCGGAGTACTGTCCTCCGCGAATTCCGGAGTACTGTCCTCCGTAGACGCTAGC |
| Enhancer sites | n = 5 x UAS binding sites | ATCGATAGGTACCGAGTTTCTAGACGGAGTACTGTCCTCCGAGCGGAGTACTGTCCTCCGACTCGAGCGGAGTACTGTCCTCCGATCGGAGTACTGTCCTCCGCGAATTCCGGAGTACTGTCCTCCGTAGACGTAGACGCTAGC |
| Core promoter | AMLP | GGGGGGCTATAAAAGGGGGTGGGGGCGTTCGTCCTCACTCT |
| Core promoter | YB-TATA | TCTAGAGGGTATATAATGGGGGCCA |

**Table S2**. DNA sequences of tested iRFP-P2A-proBMP promoter constructs (related to **Fig. S1D**).

| Bases | Part | Part Sequence |
|---|---|---|
| *Promoter driving iRFP-P2A-proBMP4 expression* | | |
| 1..154 bp | upstream polyA signal | AATAAAATATCTTTATTTTCATTACATCTGTGTGTTGGTTTTTTGTGTGAATCGATAGTACTAACATACGCTCTCCATCAAAACAAAACGAAACAAAACAAACTAGCAAAATAGGCTGTCCCCAGTGCAAGTGCAGGTGCCAGAACATTTCTCT |
| 155.. 336 bp | 3 x UAS - **AMLP** promoter | ATCGATAGGTACCGAGTTTCTAGACGGAGTACTGTCCTCCGAGCGGAGTACTGTCCTCCGCGAATTCCGGAGTACTGTCCTCCGTAGACGCTAGC**GGGGGGCTATAAAAGGGGGTGGGGGCGTTCGTCCTCACTCT**AGATCTGCGATCTAAGTAAGCTTTGCATTCCGGTACTGTTGGTAAA |
| 337.. 345 bp | Kozak | GCCACC(ATG) |
| *Promoter driving GAVPO expression* | | |
| 1..363 bp | TRE3G promoter | CTCGAGTTTACTCCCTATCAGTGATAGAGAACGTATGAAGAGTTTACTCCCTATCAGTGATAGAGAACGTATGCAGACTTTACTCCCTATCAGTGATAGAGAACGTATAAGGAGTTTACTCCCTATCAGTGATAGAGAACGTATGACCAGTTTACTCCCTATCAGTGATAGAGAACGTATCTACAGTTTACTCCCTATCAGTGATAGAGAACGTATATCCAGTTTACTCCCTATCAGTGATAGAGAACGTATGTCGAGGTAGGCGTGTACGGTGGGCGCCTATAAAAGCAGAGCTCGTTTAGTGAACCGTCAGATCGCCTGGAGCAATTCCACAACACTTTTGTCTTATACTTGGGGAATGCC |
| 364.. 372 bp | Kozak | GCCACC(ATG) |

**Table S3**. Final promoter sequences used in LightBMP cell line engineering.

| Aminoacids | Protein name | Aminoacid sequence |
|---|---|---|
| iRFP-P2A-proBMP4 fluorescent reporter and BMP morphogen | | |
| 1-327 aa | iRFP670-NES | MARKVDLTSCDREPIHIPGSIQPCGCLLACDAQAVRITRITENAGAFFGRETP<br>RVGELLADYFGETEAHALRNALAQSSDPKRPALIFGWRDGLTGRTFDISLHR<br>HDGTSIIEFEPAAAEQADNPLRLTRQIIARTKELKSLEEMAARVPRYLQAMLG<br>YHRVMLYRFADDGSGMVIGEAKRSDLESFLGQHFPASLVPQQARLLYLKNAI<br>RVVSDSRGISSRIVPEHDASGAALDLSFAHLRSISPCHLEFLRNMGVSASMSL<br>SIIIDGTLWGLIICHHYEPRAVPMAQRVAAEMFADFLSLHFTAAHHQRNSNE<br>LALKLAGLDINK |
| 328-349 aa | GSG-P2A | GSGATNFSLLKQAGDVEENPGP |
| 350-756 aa | proBMP4 | IPGNRMLMVVLLCQVLLGGASHASLIPETGKKKVAEIQGHAGGRRSGQSHE<br>LLRDFEATLLQMFGLRRRPQPSKSAVIPDYMRDLYRLQSGEEEEEQIHSTGLE<br>YPERPASRANTVRSFHHEEHLENIPGTSENSAFRFLFNLSSIPENEVISSAELRL<br>FREQVDQGPDWERGFHRINIYEVMKPPAEVVPGHLITRLLDTRLVHHNVTR<br>WETFDVSPAVLRWTREKQPNYGLAIEVTHLHQTRTHQGQHVRISRSLPQGS<br>GNWAQLRPLLVTFGHDGRGHALTRRRRAKRSPKHHSQRARKKNKNCRRHS<br>LYVDFSDVGWNDWIVAPPGYQAFYCHGDCPFPLADHLNSTNHAIVQTLVN<br>SVNSSIPKACCVPTELSAISMLYLDEYDKVVLKNYQEMVVEGCGCR* |
| Light-inducible transactivator GAVPO | | |
| 1-509 aa | GAVPO-<br>flag-tag | MAKLLSSIEQACDICRLKKLKCSKEKPKCAKCLKNNWECRYSPKTKRSPLTRA<br>HLTEVESRLERLERSIATRSHTLYAPGGYDIMGYLIQIMKRPNPQVELGPVDTS<br>VALILCDLKQKDTPIVYASEAFLYMTGYSNAEVLGRNCRFLQSPDGMVKPKS<br>TRKYVDSNTINTMRKAIDRNAEVQVEVVNFKKNGQRFVNFLTMIPVRDET<br>GEYRYSMGFQCETELQYPYDVPDYAEFQYLPDTDDRHRIEEKRKRTYETFKSI<br>MKKSPFSGPTDPRPPPRRIAVPSRSSASVPKPAPQPYPFTSSLSTINYDEFPT<br>MVFPSGQISQASALAPAPPQVLPQAPAPAPAPAMVSALAQAPAPVPVLAP<br>GPPQAVAPPAPKPTQAGEGTLSEALLQLQFDDEDLGALLGNSTDPAVFTDLA<br>SVDNSEFQQLLNQGIPVAPHTTEPMLMEYPEAITRLVTGAQRPPDPAPAPL<br>GAPGLPNGLLSGDEDFSSIADMDFSALLSQISSDYKDDDDK* |

**Table S4**. Amino acid sequences of proteins used in this study.

| Antibody | Dilution | Vendor | RRID |
|---|---|---|---|
| Goat polyclonal anti-BRACHYURY | 1:400 | R&D Systems | AB_2200235 |
| Rabbit monoclonal anti-E-CAD | 1:300 | CST | AB_2291471 |
| Mouse monoclonal anti-FLAG-TAG | 1:400 | Invitrogen | AB_1957945 |
| Rabbit monoclonal anti-FOXA2 | 1:400 | Abcam | AB_11157157 |
| Sheep polyclonal anti-FOXC2 | 1:100 | R&D Systems | AB_10973139 |
| Goat polyclonal anti-FOXF1 | 1:200 | R&D Systems | AB_2105588 |
| Mouse monoclonal anti-GATA3 | 1:50 | Santa Cruz | AB_627666 |
| Goat polyclonal anti-HAND1 | 1:300 | R&D Systems | AB_2115853 |
| Mouse monoclonal anti-ISL1 | 1:100 | DSHB | AB_2314683 |
| Rabbit monoclonal anti-LEF1 | 1:400 | CST | AB_823558 |
| Rabbit polyclonal anti-MIXL1 | 1:300 | ABClonal | AB_2770366 |
| Goat polyclonal anti-NANOG | 1:200 | R&D Systems | AB_355097 |
| Mouse monoclonal anti-N-CAD | 1:300 | Sigma-Aldrich | AB_258801 |
| Rabbit monoclonal anti-NKX2.5 | 1:400 | CST | AB_2797667 |
| Mouse monoclonal anti-OCT3/4 | 1:300 | BD Biosciences | AB_398737 |
| Rabbit monoclonal anti-pSMAD1/5/8 | 1:400 | CST | AB_2493181 |
| Goat polyclonal anti-hSNAIL | 1:200 | R&D Systems | AB_2191738 |
| Rabbit monoclonal anti-SOX2 | 1:300 | CST | AB_2714146 |
| Goat polyclonal anti-SOX17 | 1:200 | R&D Systems | AB_355060 |
| Goat polyclonal anti-TBX6 | 1:300 | R&D Systems | AB_2200834 |
| Alexa Fluor 405 donkey anti-Rabbit | 1:200 | Abcam | AB_2923541 |
| Alexa Fluor 488 donkey anti-Rabbit | 1:500 | Life Technologies | AB_2535792 |
| Alexa Fluor 488 donkey anti-Mouse | 1:500 | Life Technologies | AB_141607 |
| Alexa Fluor 488 donkey anti-Goat | 1:500 | Life Technologies | AB_2534102 |
| Alexa Fluor 555 donkey anti-Rabbit | 1:500 | Life Technologies | AB_162543 |
| Alexa Fluor 555 donkey anti-Mouse | 1:500 | Life Technologies | AB_2536180 |
| Alexa Fluor 555 donkey anti-Goat | 1:500 | Life Technologies | AB_2535853 |
| Alexa Fluor 647 donkey anti-Rabbit | 1:500 | Life Technologies | AB_2536183 |
| Alexa Fluor 647 donkey anti-Mouse | 1:500 | Life Technologies | AB_162542 |
| Alexa Fluor 647 donkey anti-Goat | 1:500 | Life Technologies | AB_2535864 |

**Table S5**. Antibodies and dilutions that were used in the study.
